# Structure-Constrained Intrinsic Timescales Across Tasks

**DOI:** 10.64898/2026.08.26.747109

**Authors:** Kaichao Wu, Raul de Palma Aristides, Ruben Herzog, Claudio Mirasso, Pierpaolo Sorrentino, Leonardo L. Gollo

## Abstract

Intrinsic neural timescale (INT) quantifies the persistence of spontaneous neural dynamics and offers a principled metric for characterizing brain-wide temporal organization. Although a hierarchy of INTs has been established during rest, how task engagement reconfigures this organization and how it is constrained by the structural connectome (SC) remain poorly understood. Here, we systematically mapped whole-brain INT using high-resolution fMRI data from the Human Connectome Project during rest and seven tasks spanning working memory, gambling, motor, language, social, relational, and emotion domains. Task engagement induced robust, regionally heterogeneous changes in INT while largely preserving the brain-wide temporal hierarchy across cognitive states. SC–INT coupling remained strong but consistently decreased during tasks, indicating that anatomical architecture continues to constrain INT, although its influence is attenuated under task demands. To investigate these findings mechanistically, we employed a multiscale, whole-brain neuronal-network model, which revealed that INT increase and peak within a broad critical-like regime. Strong SC–INT coupling, as observed empirically, emerged in the subcritical regime, weakened progressively with increasing network excitability, and reversed in the super-critical regime. These results demonstrate that task engagement reconfigures INTs while maintaining their hierarchical organization, suggesting that both resting and task states operate largely within a common subcritical dynamical regime.

## Introduction

Brain regions differ markedly in the temporal scales over which they integrate and maintain information, giving rise to a hierarchical organization of neural dynamics ^1,2^. Intrinsic neural timescale (INT) provides a direct quantitative metric for these distinct regional persistence profiles ^3–6^. This measure shows marked heterogeneity across brain regions, with sensory areas generally characterized by shorter timescales and higher-order association areas by longer timescales ^3,5,7,8^. Such regional variation is not random, but follows large-scale cortical hierarchies and structural gradients ^9–11^, including core–periphery organization ^12,13^, myeloarchitectonic variation ^14^, and macroscale functional hierarchy ^7,15,16^. This systematic increase in timescales from sensory to higher-order association regions gives rise to a hierarchy of intrinsic neural timescales, which has been proposed to support progressively longer temporal integration windows along the cortical hierarchy ^5,17–19^. Extensive evidence from studies using electrophysiology and fMRI data has shown that the hierarchy of intrinsic neural timescales is robust across modalities and species under task-free (i.e., resting state) conditions ^15,20–23^. These resting state patterns are spatially organized along cortical hierarchies and large-scale functional systems ^15,18,24^, suggesting that temporal integration is not arbitrary but constrained by the underlying structural and functional architecture of the brain ^25^.

However, cognition is inherently dynamic, requiring the brain to flexibly reconfigure neural processing in response to changing task demands. Previous work suggests that such flexibility is supported by metastable brain dynamics operating near a critical regime, enabling adaptive transitions between functional states during cognition ^26–29^. Consistent with this view, the temporal organization of brain activity, including slow intrinsic dynamics, has been proposed to provide an essential mechanistic basis for task-related cognitive processing ^29^.

Recent neuroimaging studies provide growing evidence that INTs are modulated across a wide range of cognitive contexts including visual attention ^30,31^, reward processing ^5^, mnemonic discrimination ^18^, self-related cognition ^32^, narrative construction ^33^, consciousness ^34,35^, music listening ^36^, and continuous behavior ^22^. These studies indicate that INT is not merely a static property of spontaneous activity, and can be modulated by ongoing cognitive demands and behavioral performance ^7,18^. Yet, it remains unclear to what extent such task-related changes reflect a genuine reorganization of the brain’s temporal hierarchy, or whether they remain essentially constrained by the underlying anatomical structure. Addressing this issue is essential for improving our understanding of how a fixed anatomical architecture can support a stable temporal hierarchical backbone without sacrificing the flexibility needed to meet diverse cognitive demands at the regional level.

Here, we systematically characterize whole-brain INT across seven cognitive tasks assessed using data from the Human Connectome Project (HCP) and compare the resulting patterns with those observed at rest. The tasks comprise Working Memory, Gambling, Motor, Language, Social, Relational, and Emotion (Table 1). We examine both global and regional task-related changes in intrinsic timescales, asking whether task engagement preserves the relative ordering of timescales across regions or yields task-specific deviations from the resting hierarchy. Crucially, we test the extent to which task-related INT patterns remain predictable from the structural connectome, thereby directly assessing structure–function constraints on temporal dynamics across cognitive states. To mechanistically explain the phenomena, we further develop a criticality-based approach and extend it to a multiscale model, in which intrinsic timescales, owing to critical slowing down, peak at criticality and vary systematically with the distance from it ^18,37–39^. By linking empirical task-dependent INT modulation to shifts in distance to criticality, we assess whether diverse cognitive states can be understood as structured departures from a common critical regime, jointly sculpted by anatomical connectivity and task demands.

**Table 1.**
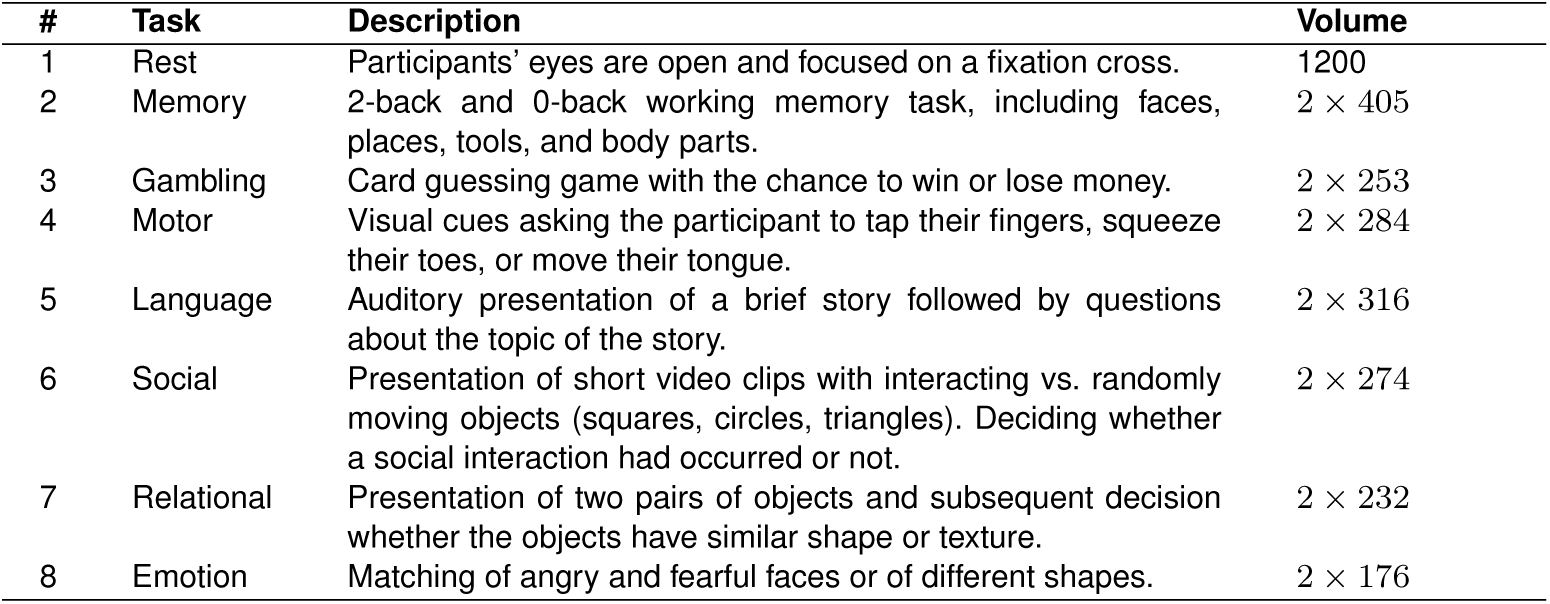
Overview of HCP fMRI conditions included in the study.

## Results

We investigated how intrinsic neural timescales are organized by brain structure and dynamically reconfigured across cognitive states. Resting-state and task fMRI data spanning multiple cognitive domains from 326 neurologically intact adults were obtained (Figure 1**A**). Diffusion MRI for each participant was utilized to estimate interregional white-matter pathways (Figure 1**B**), and then to identify structural connectomes (Figure 1**C**). We exploited the BOLD timeseries from fMRI and structural connectome to examine the interactions among anatomical connectivity, neural timescales, and large-scale brain dynamics. With the extracted BOLD timeseries at the regional level Figure 1**D**, INT was calculated by integrating the area under the curve of the autocorrelation function (ACF) from the first lag to the first zero-crossing Figure 1**E**. Relying on estimated short- and long-INTs across task conditions, we mapped task-related intrinsic timescales and their hierarchical structure. Our main question was whether the temporal hierarchy of intrinsic timescales recapitulates the underlying spatial organization.

**Fig. 1.**
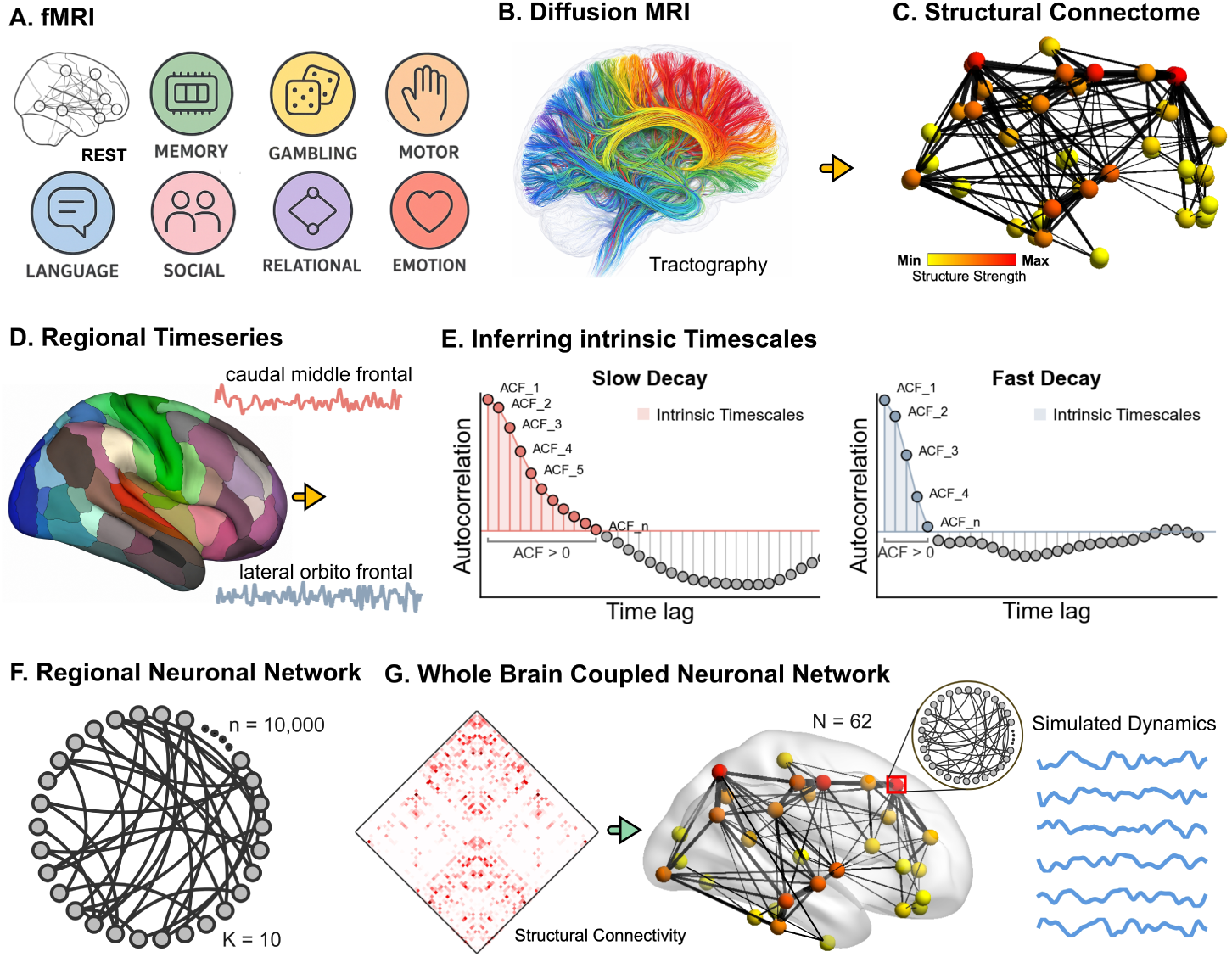
Inferring intrinsic neural timescales with BOLD timeseries and modeling dynamics. **(A)**: Resting-state and task fMRI data were acquired from the HCP dataset across participants. **(B)**: White matter tractography was obtained with the diffusion MRI for each participant, and **(C)**, structural connectomes were constructed through the tractography to define the anatomical backbone of large-scale brain networks. The colors indicate the node strength of brain regions derived from the structural connectivity matrix. **(D)**: Regional BOLD timeseries were extracted with the given Desikan–Killiany brain parcellation. Two examples of timeseries of lateral orbitofrontal (sky blue) and caudal middle frontal (pink) are from the first HCP participant. **(E)**: Intrinsic timescales (INT) were quantified from regional BOLD timeseries using the decay of the autocorrelation function, capturing the temporal integration properties of local neural activity. Different regions exhibit different decays, leading to a diversity of intrinsic timescales across the brain. As demonstrated in the illustrative plot, the lateral orbitofrontal in the left hemisphere shows a slow-decaying ACF with shorter INTs (sky blue ACF plot), whereas the caudal middle frontal in the left hemisphere shows a fast-decaying ACF (pink ACF plot). To further examine intrinsic timescales, neuronal networks were implemented at multiple spatial scales: **(F)**, a single regional neuronal network, and **(G)**, a whole-brain coupled neuronal network, where multiple regional neuronal networks were coupled based on the structural connectome, and dynamical and computational properties were obtained with two models.

To mechanistically understand the interplay between anatomical connectivity and INTs, a multiscale structure-constrained whole-brain computational neuronal network model was implemented. Each brain region was modeled as a large network of excitable units using the Kinouchi-Copelli model ^37^ (Figure 1**F**). These regional networks were then coupled using inter-regional connections derived from the estimated structural connectivity (Figure 1**G**), and regional dynamics were characterized by extracting the timeseries of the instantaneous density of active nodes within each brain region. These signals were subsequently used to compute the intrinsic timescales of the simulated brain activity.

### Intrinsic neural timescales and their hierarchy across rest and multiple tasks

With denoised BOLD timeseries of all task conditions, INT was computed for the resting-state and multiple tasks. The maps of INT are shown in Figure 2**A**, indicating that different cognitive tasks generate distinct spatial patterns of intrinsic timescales. To further visualize the global trend of INT, the different conditions were ordered according to their mean INT values Figure 2**B**. This ordering revealed a gradient from Social (*Mean ± Std*: 2.91 *±* 0.46), Relational (*Mean ± Std*: 2.65 *±* 0.39), Emotion (*Mean ± Std*: 2.62 *±* 0.43), Motor (*Mean±Std*: 2.58*±*0.34), Gambling (*Mean±Std*: 2.54*±*0.45), Rest (*Mean±Std*: 2.44 *±* 0.32), and Memory (*Mean ± Std*: 2.43 *±* 0.34) to Language (*Mean ± Std*: 2.38 *±* 0.34). A repeated-measures ANOVA reveals a significant main effect of task on the subject-specific INT, *F* (7, 326) = 90.3105, Greenhouse-Geisser (GG) corrected p-value *p_GG_ <* 0.0001, demonstrating that intrinsic timescales were significantly modulated by task demands.

**Fig. 2.**
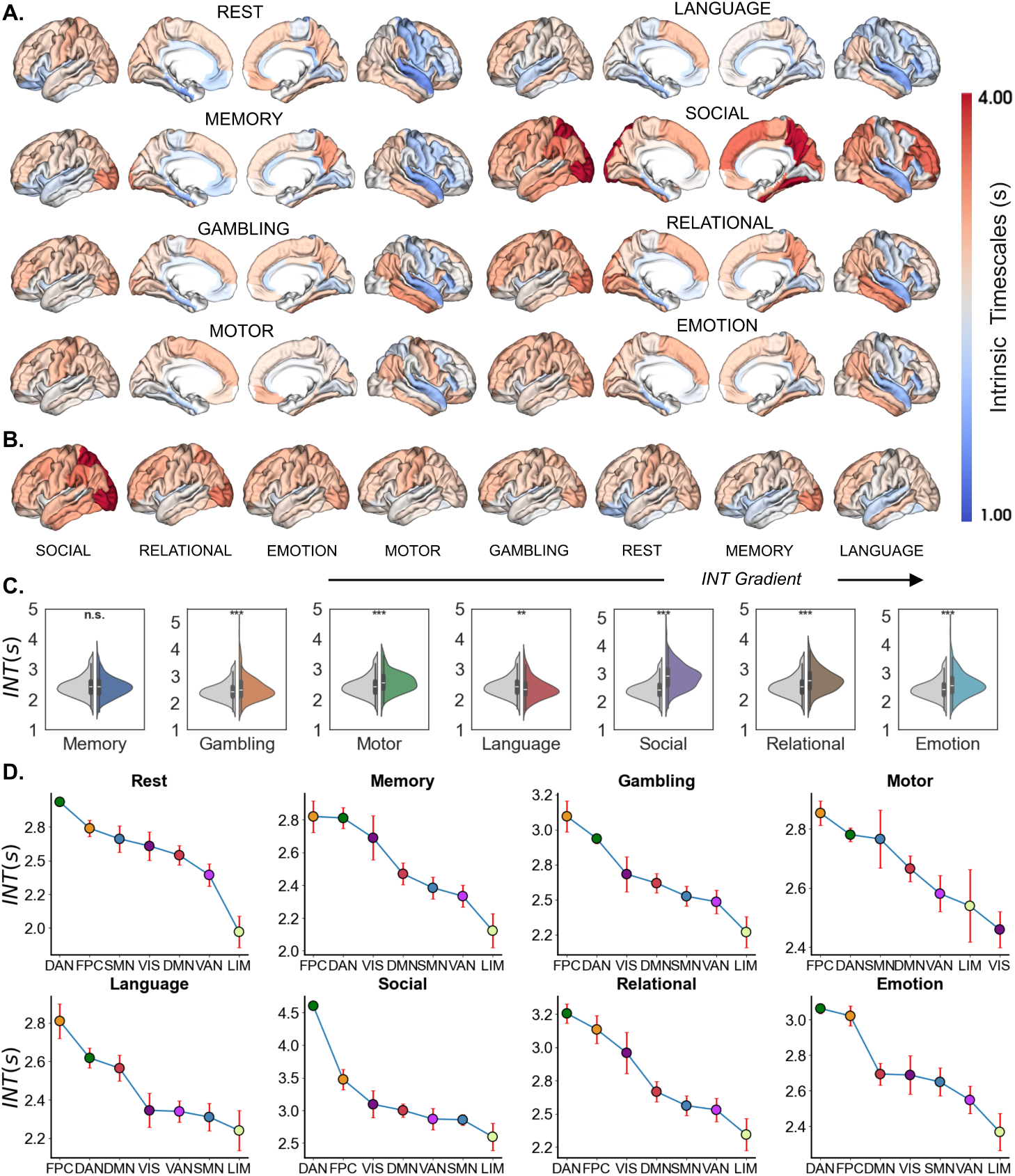
Intrinsic timescales and temporal hierarchy across rest and task states. **(A)**: The INT map across rest and 7 tasks, Memory, Gambling, Motor, Language, Social, Relational, and Emotion. **(B)**: Global mean intrinsic timescales across resting state and task conditions. **(C)**: A repeated-measures ANOVA revealed a significant main effect of task on the subject-specific INTs, *F* (7, 326) = 90.3105, Greenhouse-Geisser (GG) corrected p-value *p_GG_ <* 0.0001. Compared with rest, INT decreased during Memory (*t* = *−*0.75, *p_bhF DR_* = 0.4499) and Language (*t* = *−*3.088, *p_bhF DR_* = 0.0029), whereas Gambling (*t* = 3.778, *p_bhF DR_* = 0.0002), Motor (*t* = 6.0939, *p_bhF DR_ <* 0.0001), Social (*t* = 16.0819, *p_bhF DR_ <* 0.0001), Relational (*t* = 8.0591, *p_bhF DR_ <* 0.0001) and Emotion tasks (*t* = 6.6430, *p_bhF DR_ <* 0.0001) elicited significant INT increases. Asterisks indicate statistical significance: \**p <* 0.05, \*\**p <* 0.01, \*\*\**p <* 0.001, and \*\*\*\**p <* 0.0001 and *ns* denotes no significance detected. **(D)**: Hierarchy of intrinsic timescales across functional networks: dorsal attention (DAN), frontoparietal control (FPC), somatomotor (SMN), visual (VIS), default (DMN), ventral attention (VAN), and limbic networks (LIM). The correlation between the hierarchy rank of task and rest ranged from *r* = *−*0.42 to *r* = 0.10 with all comparisons being non-significant (*p_F DR_ >* 0.33), demonstrating that the brain temporal hierarchy was preserved across tasks.

As a crucial benchmark, the resting state captures endogenous, task-independent brain organization and offers a stable reference for quantifying task-specific reorganizations. To this end, we compared INT across task conditions against those measured during rest. Post-hoc pairwise comparisons, conducted using paired-samples t-tests with Benjamini–Hochberg false discovery rate (bhFDR) correction, revealed robust and systematic INT reorganization across most task conditions relative to rest. Specifically, INT significantly decreased during the Language task (*t* = –3.088, *p_bhF DR_* = 0.0029), whereas significant increases were observed for Gambling (*t* = 3.778, *p_bhF DR_* = 0.0002), Motor (*t* = 6.0939, *p_bhF DR_ <* 0.0001), Social (*t* = 16.0819, *p_bhF DR_ <* 0.0001), Relational (*t* = 8.0591, *p_bhF DR_ <* 0.0001), and Emotion tasks (*t* = 6.6430, *p_bhF DR_ <* 0.0001), see Figure 2**C**. Although the global mean INT did not differ significantly for the Memory condition (*t* = –0.75, *p_bhF DR_* = 0.4499), widespread regional differences between this task and rest were evident (Supplementary Materials Figure **1**).

To characterize the temporal hierarchy of intrinsic timescales ^19,40^, we aggregated the cortical INT within seven large-scale functional networks obtained from the Yeo et al. ^41^: dorsal attention (DAN), frontoparietal control (FPC), somatomotor (SMN), visual (VIS), default (DMN), ventral attention (VAN), and limbic networks (LIM). The hierarchy of intrinsic timescales at this large-scale functional network level across tasks is shown in Figure 2**D**, with sensory systems tending to show shorter INT and association networks exhibiting longer INT. Each task-specific rank of network-level INT was then compared with the rank of resting-state using Spearman’s rank correlation. As a result, the correlation coefficients ranged from *r* = *−*0.42 to *r* = 0.11 (Memory: *r* = *−*0.428, *p* = 0.3373; Gambling: *r* = *−*0.428, *p* = 0.3373; Motor: *r* = *−*0.428, *p* = 0.3373; Language: *r* = 0.001, *p* = 0.9972; Social: *r* = 0.1071, *p* = 0.8191; Relational: *r* = *−*0.1071, *p* = 0.8191; Emotion: *r* = 0.1071, *p* = 0.8191) with all comparisons being non-significant (*p_F DR_ >* 0.33). These weak correlations, together with the absence of significant changes in the ranking of functional networks relative to the resting state, indicate that the overall hierarchical organization of intrinsic timescales was largely preserved across task conditions. Notably, the Memory and Gambling tasks exhibited identical INT rankings across functional networks.

The largely consistent hierarchy of INT across different task conditions suggests that INT reflects a predominantly stable and general principle of brain functional organization, even when the brain responds to different stimuli, rather than a strongly modulated task-specific temporal pattern. As shown in the Supplementary Materials Figure **2**, this largely preserved hierarchy of INTs is also observed in the Glasser360 parcellation ^42^.

### Task-related changes in intrinsic timescales are associated with cognitive functional maps

To examine the cognitive relevance of task-induced intrinsic timescales, we computed regional INT difference maps for each task condition relative to resting state (Figure 3**A-G**). These maps revealed spatially heterogeneous patterns of INT changes across the cortex, indicating that tasks induced distinct regional deviations in timescales from the resting state. We then assessed the dissimilarity between task-induced INT modulation patterns across tasks by calculating the pairwise distance between task difference maps, defined as 1 *− r*, where *r* denotes the spatial correlation between two maps. The distance matrix in Figure 3**H** shows that the mean distances ranged from 0.54 to 0.85, with the smallest distance between Memory and Gambling (*Mean ± Std* : 0.54 *±* 0.08) and the largest between Social and Motor (*Mean ± Std* : 0.85 *±* 0.13). A similar behavior is also observed for the Glasser360 parcellation (Supplementary Materials Figure **3**). Hence, as expected, different tasks require distinct magnitudes and spatial configurations of INT modulation, suggesting that distinct cognitive demands are supported by correspondingly distinct patterns of intrinsic timescale regulation.

**Fig. 3.**
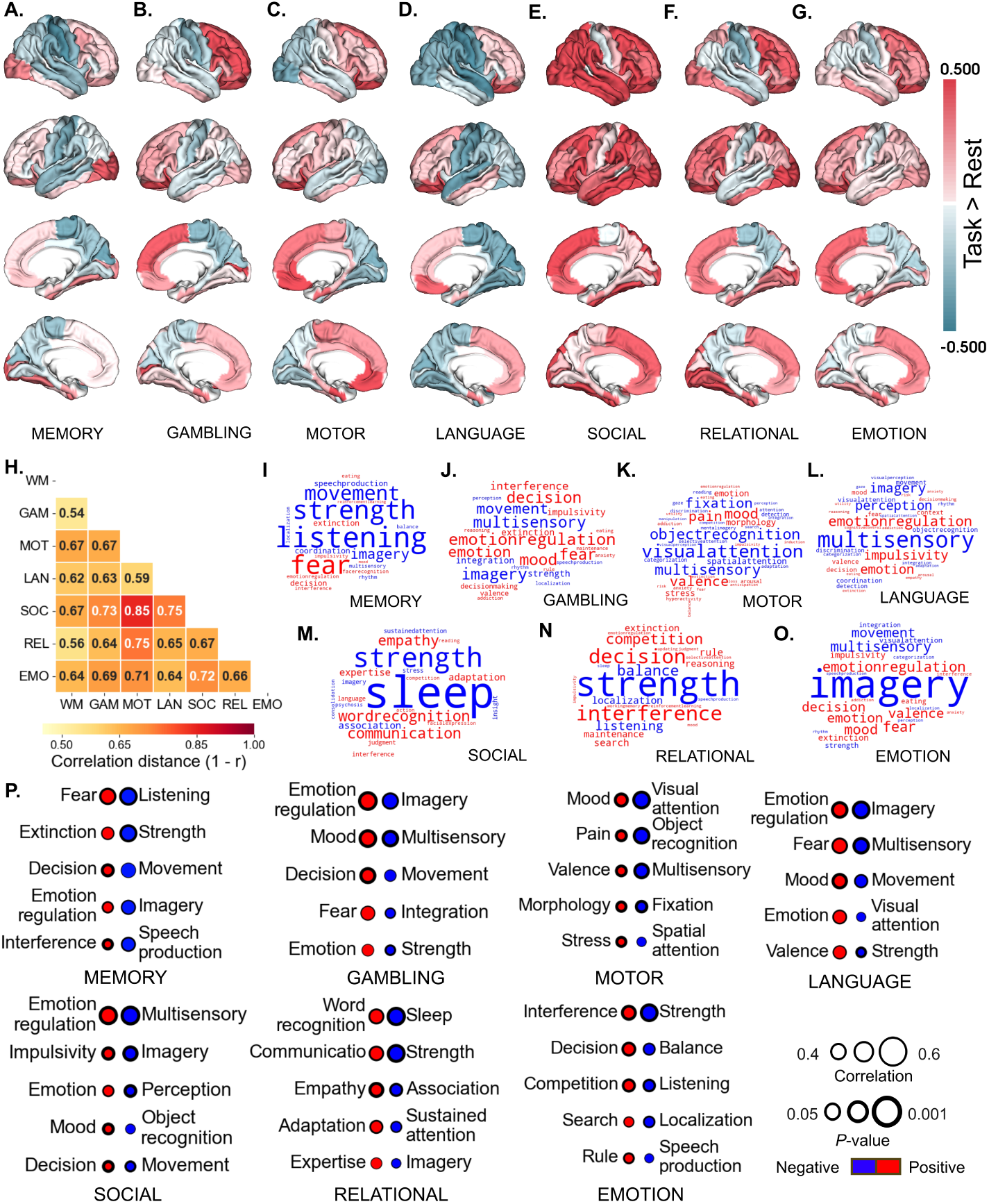
Task-induced changes in INT and their cognitive associations. **A-G**: Regional INT difference maps for the Memory **(A)**, Gambling **(B)**, Motor **(C)**, Language **(D)**, Social **(E)**, Relational **(F)**, and Emotion **(G)** tasks. Warmer colors indicate regions with higher INT during task relative to rest, whereas cooler colors indicate regions with lower INT during task relative to rest. **H**: Pairwise spatial distance between task-specific INT difference maps. Working Memory (WM), Gambling (GAM), Motor (MOT), Language (LAN), Social (SOC), Relational (REL), and Emotion (EMO). The color bar indicates correlation distance (1 *− r*), with lighter colors representing smaller distances and more similar INT difference maps, and darker red colors representing larger distances and more distinct INT difference maps. **I–O**: Neurosynth-based cognitive decoding results for the corresponding task-related INT difference maps. Word clouds show 123 cognitive terms whose meta-analytic spatial maps were significantly associated with each task difference INT map. Red and blue terms indicate positive and negative spatial associations, respectively, and word size reflects the strength of the correlation. **P**: The top 5 positive and negative associated words and the FDR-corrected p-values are shown below. The thickness of the circles represents statistical significance. The p-values were computed using the Spin test up to 10,000 permutations before performing the FDR correction.

We next decoded these task difference maps using Neurosynth meta-analytic cognitive maps ^43^. The Neurosynth meta-analytic engine provides functional activation maps associated with 123 cognitive and behavioral terms derived from the Cognitive Atlas ^44^. For each task, spatial correlations between the INT difference map and Neurosynth maps were used to identify cognitive terms whose cortical distributions were most strongly associated with task-related INT changes (see detailed description in Methods Neurosynth Association). The resulting decoding profiles revealed interpretable and task-specific cognitive associations Figure 3**I-P**, reflecting the expected characteristics of each task domain, including working memory, reward processing, emotion recognition, motor execution, language comprehension, social inference, and relational reasoning(see the detailed task battery description in Table 1 of Methods The HCP battery of seven tasks). In general, INT modulation during task states captures both task-specific cognitive demands and shared processes such as attention, decision-making, imagery, affective processing, and response selection.

### Brain structure constrains intrinsic timescales across resting state and tasks

The brain’s structural wiring has been viewed as scaffolding its functional dynamic activity ^45–49^. To investigate how the structural architecture shapes task-dependent brain dynamics, we examine the relationship between regional SC strength (see Figure 4**A**&**C** for two atlases) and INT. This measure, termed SC-INT coupling as normally indicated in previous literature on structural and functional connectome ^47,50,51^, was computed as the Pearson correlation between regional INT values and regional structural connectivity strength. We found that SC-INT coupling was significantly positive across all conditions (see Figure 4**B**&**D**), suggesting that the temporal hierarchy of brain dynamics remains strongly shaped by the underlying structural connectome regardless of the cognitive demands.

**Fig. 4.**
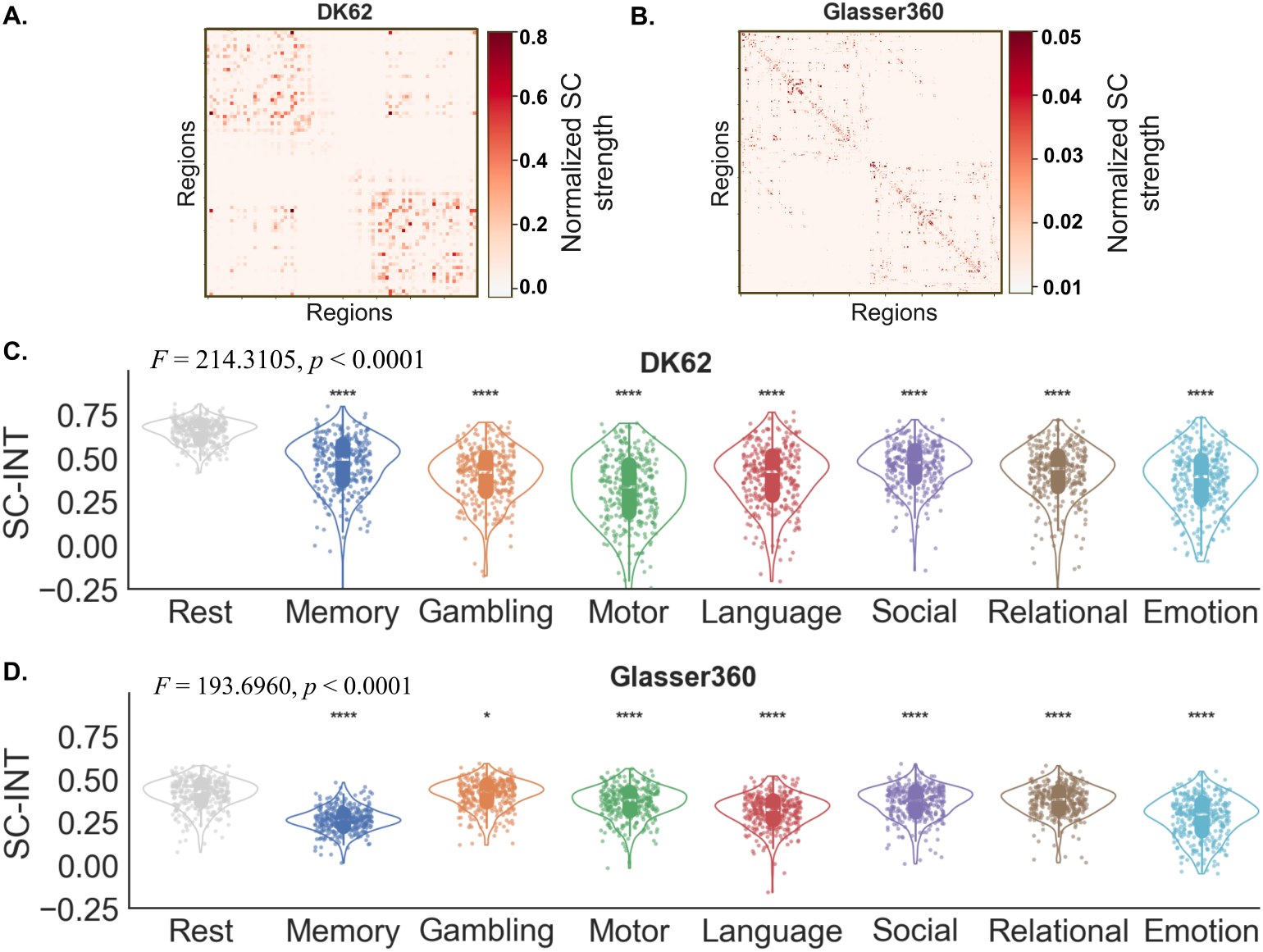
Structural connectivity constrains intrinsic timescales across rest and task states. **(A)**: Normalized structural connectivity matrix for DK62. **(B)**: Normalized structural connectivity matrix for Glasser360. **(C)**: SC-INT coupling, quantified as the Pearson correlation between regional intrinsic timescale (INT) values and regional SC strength, for DK62. Each point represents one subject. A repeated-measures ANOVA shows a significant task effect on SC-INT coupling DK atlas: *F* (7, 2282) = 214.1051, *p_GG_ <* 0.0001. **(D)** Same for the Glasser360 atlas: *F* (7, 2282) = 193.6960, *p_GG_ <* 0.0001), and post hoc pairwise test indicated a weakened SC-INT coupling during all tasks relative to rest (*p_bhF DR_ <* 0.01). . Asterisks indicate statistical significance relative to rest: \**p <* 0.05, \*\**p <* 0.01, \*\*\**p <* 0.001, and \*\*\*\**p <* 0.0001.

A repeated-measures ANOVA (rmANOVA) on subject-level coupling values revealed a significant main effect of task (*F* (7, 2282) = 214.1051, *p_GG_ <* 0.0001), indicating that the strength of SC-INT coupling varied across conditions (Figure 4**C**). Post-hoc paired comparisons further showed that SC-INT coupling was significantly weaker in all task conditions than in the resting state (all *p_F DR_ <* 0.0001). Among these tasks, Motor showed the largest difference relative to rest (*t* = *−*34.59, *p_F DR_ <* 0.0001), followed by Gambling (*t* = *−*32.71, *p_F DR_ <* 0.0001), Emotion (*t* = *−*31.60, *p_F DR_ <* 0.0001), Relational (*t* = *−*30.87, *p_F DR_ <* 0.0001), Language (*t* = *−*30.27, *p_F DR_ <* 0.0001), Social (*t* = 28.93, *p_F DR_ <* 0.0001), and Memory (*t* = *−*24.16, *p_F DR_ <* 0.0001). The same result can be observed in fine-grained 360 parcellation (Figure 4**D**), where SC-INT coupling was significantly different across tasks (rmANOVA with GG multiple correction; *F* (7, 2282) = 193.6960, *p_GG_ <* 0.0001) and was weakened in task conditions; Memory: *t* = *−*28.66, *p_F DR_ <* 0.0001); Gambling: *t* = *−*2.28, *p_F DR_ <* 0.01); Motor: *t* = *−*8.43, *p_F DR_ <* 0.0001); Language: *t* = *−*16.52, *p_F DR_ <* 0.0001); Social: *t* = *−*8.27, *p_F DR_ <* 0.0001); Relational: *t* = *−*7.14, *p_F DR_ <* 0.0001). The spatial correlation between SC and INT further showed that task-based INT–SC coupling was consistently and strongly positive across tasks, ranging from *r* = 0.60 to *r* = 0.74 for DK62 and from *r* = 0.34 to *r* = 0.61 for Glasser360 (Figure 5A&B). For all cases, these correlations were highly significant after correction for multiple comparisons (*p_F DR_ <* 0.0001).

**Fig. 5.**
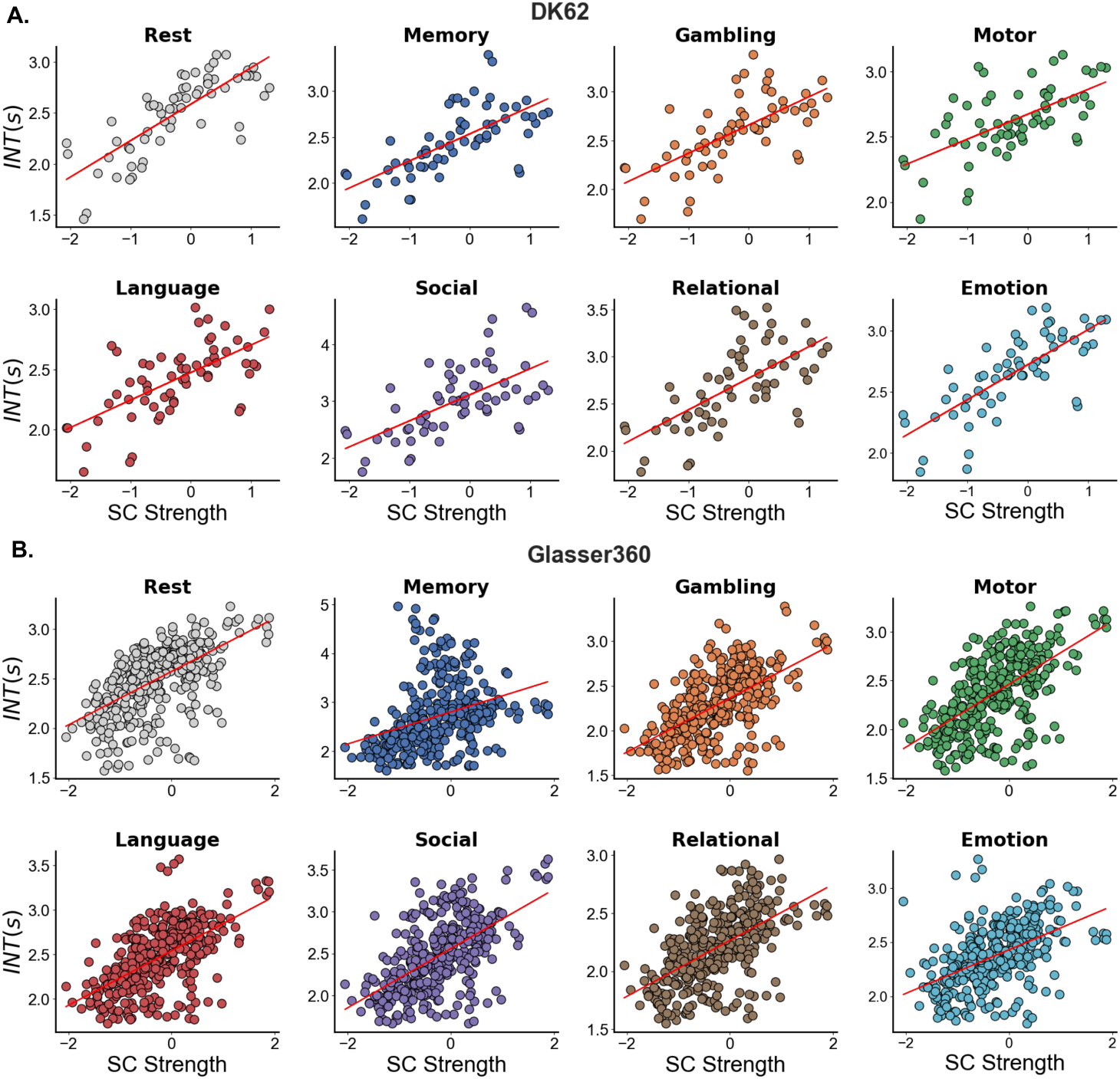
Spatial correlations between structural connectivity and intrinsic timescale across task states. Scatter plots show the node-wise association between SC and INT for each task condition using the DK62 atlas **(A)** and Glasser360 atlas **(B)**. With DK62, emotion showed the strongest node-wise coupling (*r* = 0.74, *p_F DR_ <* 0.0001), followed by Memory (*r* = 0.68, *p_F DR_ <* 0.0001), Gambling (*r* = 0.67, *p_F DR_ <* 0.0001), Relational (*r* = 0.66, *p_F DR_ <* 0.0001), Language (*r* = 0.64, *p_F DR_ <* 0.0001), Social (*r* = 0.62, *p_F DR_ <* 0.0001), and Motor (*r* = 0.60, *p_F DR_ <* 0.0001). With Glasser360, Motor showed the strongest coupling (*r* = 0.61, *p_F DR_ <* 0.0001), followed by Language (*r* = 0.59, *p_F DR_ <* 0.0001), Gambling (*r* = 0.59, *p_F DR_ <* 0.0001), Social (*r* = 0.58, *p_F DR_ <* 0.0001), Relational (*r* = 0.56, *p_F DR_ <* 0.0001), Emotion (*r* = 0.50, *p_F DR_ <* 0.0001), and Memory (*r* = 0.34, *p_F DR_ <* 0.0001). Each dot represents a regional value averaged across subjects.

### Critical slowing down emerges in a multiscale brain model

To elucidate how the structural connectome shapes the hierarchy of intrinsic timescales, we developed a multiscale neuronal network model that integrates local circuit interactions within brain regions with long-range interactions mediated by the whole-brain connectome (Figure 1F–G, See Methods Neuronal network modelling). Each brain region was modeled as a single regional neuronal network with *n* = 10,000 neurons and an average degree *K* = 10. Based on diffusion MRI tractography, the structural connectome and the structural connectivity matrix (Figure 4 A&C) were obtained and then used to couple multiple regional neuronal networks into a whole-brain neuronal network (Figure 1G).

Both the single regional networks and the coupled whole-brain network employed the Kinouchi–Copelli dynamics model (described in Methods: Neuronal Dynamics) to generate simulated brain activity Figure 1G), from which INT were computed using the autocorrelation-based approach (see Figure 1E). At the regional level, intrinsic timescales exhibited a strong dependence on the branching ratio *σ* (Figure 6A, blue curve). In isolated regional networks, INT increased markedly as *σ* approached the critical point, reaching a maximum at *σ* = 1, consistent with the phenomenon of critical slowing down ^37,52–55^. When regions were coupled through the whole-brain structural connectome, the global average INT likewise peaked near *σ_g_*= 1 (Figure 6A, red curve).

**Fig. 6.**
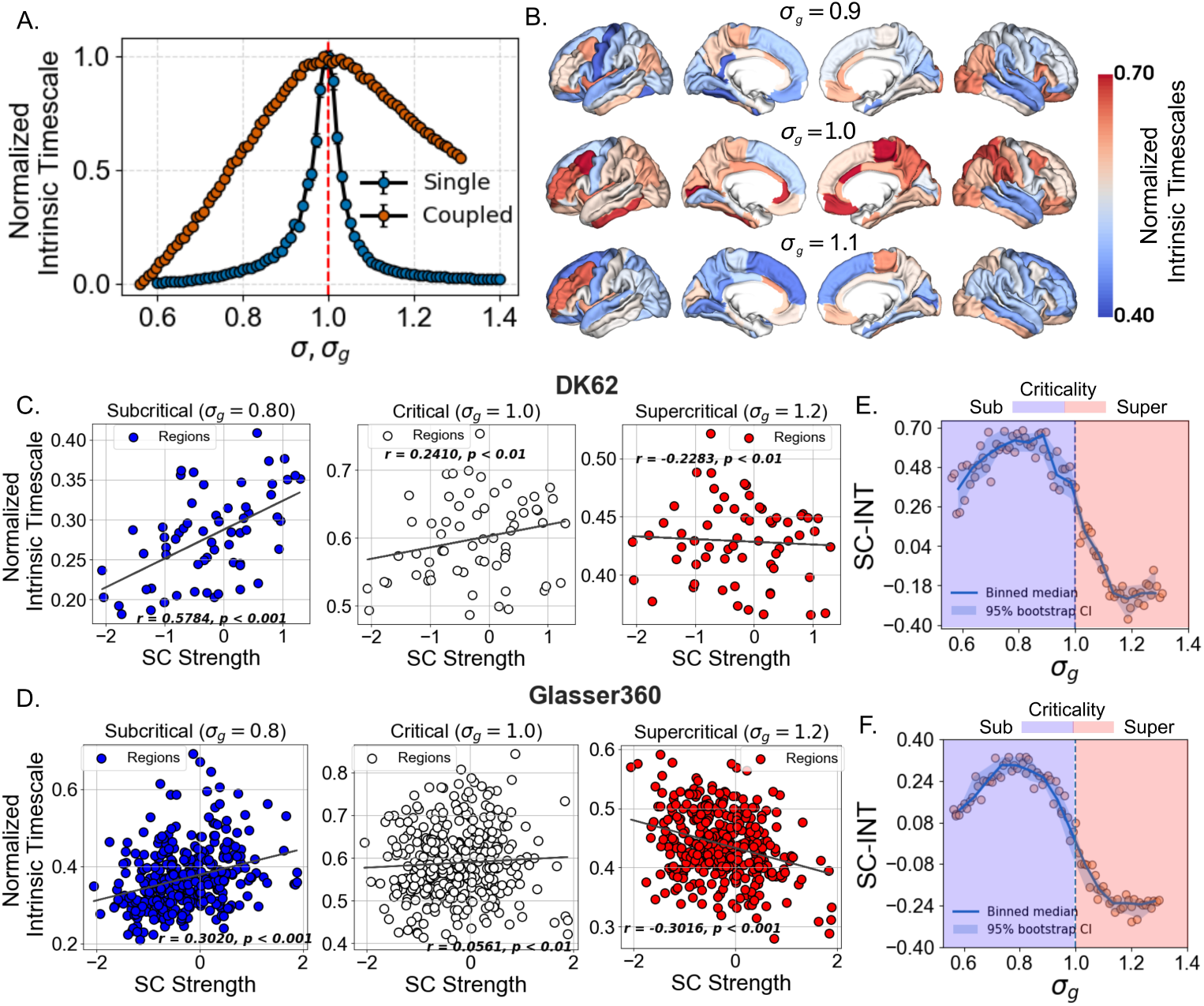
Modeling reveals intrinsic timescales and their relationship with structural connectivity across dynamical regimes. **(A)**: Intrinsic timescales as a function of the control parameter *σ* for single (blue) and *σg* for coupled (red) networks. In the isolated single network, intrinsic timescales exhibit a sharp peak at *σ ≈* 1. The coupled network also peaks at *σg ≈* 1, whereas it shows a broader peak, indicating an extended regime of elevated timescales. After normalization, the coupled network displays a markedly broader profile compared to the single network, highlighting the widening of the near-critical regime due to coupling. **(B)**: Spatial maps of intrinsic timescales across different regimes. The color bar indicates the normalized intrinsic timescale. Scatter plots of SC strength versus intrinsic timescale for representative regimes (subcritical, critical, supercritical) for DK62 **(C)** and Glasser360 **(D)**. **(E)&(F)**: SC-INT coupling as a function of *σg* for DK62 and Glasser360 shows a clear transition. The solid line represents the binned median, and the shaded area indicates 95% bootstrap confidence intervals.

To compare the shape of these relationships, we normalized INT values across multiple scales of the neuronal network. The resulting profiles revealed a striking difference between isolated single and coupled networks (Figure 6A). In isolated networks, INT displayed a narrow and sharply defined peak around the critical point. In contrast, the whole-brain network exhibited a broader, smoother peak, suggesting that enhanced intrinsic timescales are maintained over a wider range of parameter values. These findings indicate that structural coupling broadens the critical regime, reducing the need for precise parameter tuning and increasing the robustness of enhanced timescales. For the whole-brain structural network, we obtained the spatial distribution of regional INT across different dynamical regimes (Figure 6B). These maps enable comparison with regional structural connectivity strength, providing insight into the potential mechanisms underlying the observed relationship. Similar results were also obtained for the Glasser360 parcellation (Supplementary Materials Figure **4**).

### Structural constraints on intrinsic timescales arise in the subcritical regime

To further determine how SC-INT coupling depends on the underlying dynamical regime, we examined the relationship between structural connectivity and simulated intrinsic timescales (INT) across subcritical, critical, and supercritical conditions. Using the DK62 parcellation (Figure 6C), regional INT was strongly positively correlated with structural connectivity in the subcritical regime (*σ_g_* = 0.8; *r* = 0.5784, *p <* 0.001). This association weakened substantially near criticality (*σ_g_* = 1.0; *r* = 0.2410, *p <* 0.01) and reversed in the supercritical regime (*σ_g_* = 1.2; *r* = *−*0.2283). A similar pattern was observed with the Glasser360 parcellation (Figure 6D), demonstrating that the relationship is robust across spatial resolutions.

To characterize this transition more systematically, we quantified SC-INT coupling across the full range of *σ_g_*. As shown in Figure 6E,F, SC-INT coupling remained positive throughout the subcritical regime, reaching its maximum around *σ_g_ ≈* 0.8, before progressively declining as the system approached the critical point. Near criticality, the coupling remained positive but was markedly attenuated, whereas in the supercritical regime it became negative. This smooth transition was highly consistent across both parcellation schemes.

These findings demonstrate that the influence of structural connectivity on intrinsic timescales is strongly dependent on the underlying dynamical regime. Below criticality, highly connected regions tend to exhibit longer intrinsic timescales, indicating that anatomical connectivity strongly constrains regional temporal integration. As the system approaches criticality, this structural constraint weakens, suggesting that temporal dynamics become less dependent on differences in structural connectivity. Beyond the critical point, the relationship reverses, with structurally stronger regions tending to exhibit shorter intrinsic timescales. Together, these results show that the coupling between brain structure and intrinsic timescales is not fixed but is continuously reshaped by the global dynamical state of the system.

## Discussion

This study systematically mapped intrinsic neural timescales (INTs) and their hierarchical organization across resting state and seven cognitive task conditions, while also characterizing the relationship between brain structure and temporal dynamics. Using structurally constrained dynamical models, we further investigated how anatomical connectivity shapes the temporal organization of brain activity across cognitive states. Three key findings emerged from the empirical data. First, cognitive tasks induced significant but regionally heterogeneous modulations of INTs. Second, although task engagement altered absolute INT values, the large-scale cortical hierarchy of timescales remained remarkably preserved, with longer timescales in high-order association networks and shorter timescales in unimodal networks. Third, INTs showed a consistent positive association with structural connectivity (SC) strength across all tasks, with the strongest coupling observed during rest. These findings indicate that task-related changes in neural timescales are not arbitrary, but instead occur within a structured dynamical landscape constrained by the underlying anatomical architecture.

Complementing these empirical findings, our multiscale computational modeling demonstrates that heterogeneity across brain regions broadens the critical-like regime and supports enhanced INTs ^56^, rather than producing a sharp peak as observed for a single brain region. Importantly, the model further shows that the relationship between SC and INT is strongly dependent on the underlying dynamical regime ^57^. A hierarchical organization of timescales accompanied by strong SC–INT coupling emerges when most brain regions operate in the subcritical regime. In this regime, regions with greater structural connectivity exhibit longer intrinsic timescales, consistent with critical slowing down as their branching ratios approach the critical point (*σ* = 1). As the system approaches criticality (*σ_g_* = 1), however, SC–INT coupling weakens, consistent with a regime in which temporal integration becomes broadly enhanced across the network, including conditions in which some highly connected hub regions already exhibit signatures of supercritical activity (Figure 7). When the entire network enters the supercritical regime (*σ_g_ >* 1), the relationship is reversed: structurally stronger regions tend to exhibit shorter timescales, again as a consequence of critical slowing down, but now because these regions lie farther from the critical point.

**Fig. 7.**
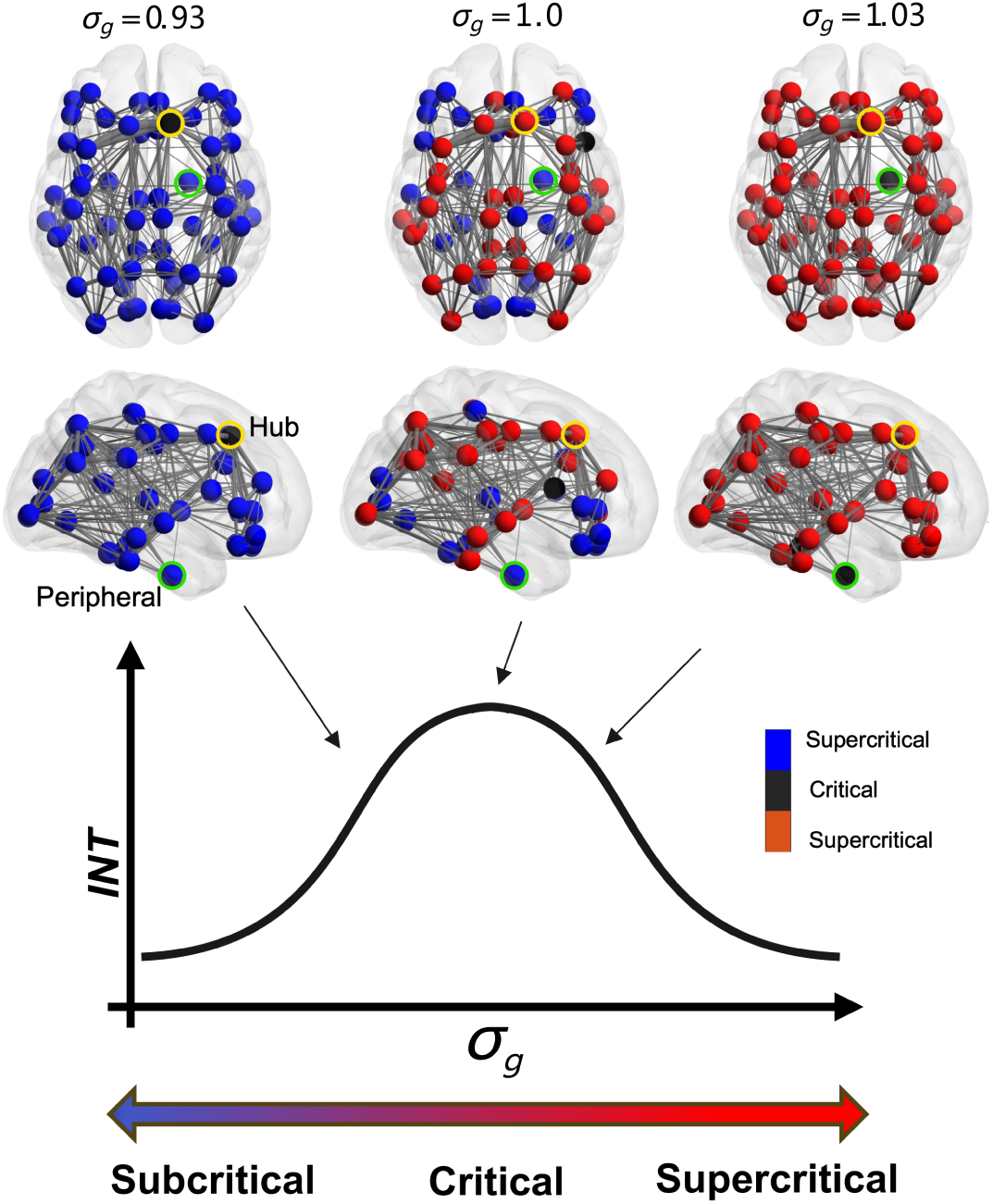
Heterogeneous distance-to-criticality in whole-brain dynamics. Sequential critical transitions occur for brain regions as the global branching ratio (*σg*) increases, with structurally well-connected hubs reaching criticality ahead of peripheral regions. In the subcritical regime, this heterogeneity gives rise to a stable hierarchy of intrinsic neural timescales, where hubs display slower fluctuations consistent with critical slowing down. Excessive increase in *σg* drives the network into the supercritical regime, which collapses this hierarchy of timescales.

The subcritical regime therefore provides the closest correspondence to the empirical observation of strong SC–INT coupling. This finding is consistent with a broader literature suggesting that near-critical subcritical dynamics can preserve many of the functional advantages associated with criticality while maintaining greater dynamical stability ^58–64^. Structural heterogeneity may further extend these advantages by allowing different regions to operate at distinct points along the dynamical landscape. As illustrated in Figure 7, regions operating in the subcritical regime may provide more stable and reliable dynamics while maintaining a buffer from the potentially pathological dynamics associated with the supercritical regime, whereas regions at or very near criticality may benefit from enhanced sensitivity and responsiveness ^38^. Such coexistence could allow the network as a whole to simultaneously exploit complementary dynamical properties, rather than requiring all regions to operate at a single optimal point. Thus, heterogeneity may not merely broaden the range of dynamical states accessible to the network, but may also provide a mechanism for distributing distinct computational advantages across its anatomical organization. Overall, our results demonstrate that the influence of anatomical connectivity on temporal dynamics is not fixed, but emerges from an interaction between network topology and dynamical state, potentially allowing the brain to simultaneously support stability, sensitivity, and flexible temporal integration.

This regime-dependent mechanism provides a potential explanation for two seemingly opposing observations: the persistence of a stable cortical hierarchy of timescales alongside substantial task-dependent variability in regional INTs. Our findings suggest that cognitive states do not fundamentally reorganize the temporal architecture of the brain. Rather, task engagement induces structured reconfigurations of neural dynamics within a relatively stable anatomical scaffold. From this perspective, the structural connectome acts as a backbone that constrains the repertoire of possible temporal states, while shifts in local and global dynamical regimes provide the flexibility required to adapt to changing cognitive demands. The preservation of the large-scale hierarchy may therefore reflect the stabilizing influence of anatomical organization, whereas regional modulation of INTs may reflect the capacity of the underlying dynamics to selectively tune temporal integration according to behavioral context. More broadly, these results support a view of cognition as emerging from the interaction between stable structural organization and flexible dynamical processes, providing a mechanistic framework for understanding how the brain can balance robustness and adaptability across behavioral states.

### Hierarchical intrinsic timescales are shaped by macroscale structural gradients

Early electrophysiological studies demonstrated that primate cortex exhibits a hierarchy of intrinsic timescales that increases from sensory regions to prefrontal cortex ^5^. Subsequent fMRI studies, either from the small ^2,24,65^ or large scale dataset ^16,66^, have shown that a consistent hierarchy of INT exists. Unimodal regions, such as sensory and motor networks, typically exhibit shorter INT, whereas transmodal regions, including higher-order networks like the central-executive network (CEN), dorsal attention network (DAN), and default-mode network (DMN), tend to display longer INT ^2,16,19,67,68^. The hierarchy of INT is also conserved across species ^69^, and is closely related to the progressive increase of information integration capacity from sensory to transmodal regions ^8,19^. Consistent with these studies, our results show that the cortical temporal hierarchy remains largely preserved across tasks, where the high-order association networks, such as DAN and FPC networks, have longer timescales, and unimodal sensory and motor networks, such as VIS and SMN networks, have shorter timescales. Note that the precise network-level hierarchy may be subject to the network definition and parcellation resolution. Coarse atlases can average across functionally heterogeneous regions, whereas finer parcellations may reveal distinct temporal properties within the same large-scale network ^41,42,70^. This is particularly relevant for distributed association networks, such as the DMN and frontoparietal systems, whose boundaries and subdivisions vary across commonly used atlases ^71,72^. Therefore, although the magnitude and ranking of network-level INT may vary with the selected parcellation scheme, the current evidence supports that the broad unimodal-to-transmodal INT gradient is maintained across tasks.

Meanwhile, macroscale spatial organization of the cortex also follows hierarchical gradients, such as the principal functional gradient extending from sensory areas to the default mode network ^72^, which is further linked to gradients in cortical microstructure, myelination, and gene expression ^66,67^.The consistent hierarchy of intrinsic timescales across tasks suggests that the spatial organization of INT is fundamentally constrained by the hierarchical architecture of the cortex rather than by transient cognitive states alone. Nevertheless, the SC-INT may not be interpreted as evidence that connection strength alone is either necessary or sufficient for generating the cortical timescale hierarchy. Recurrent interactions and the spatial organization of connectivity have been proposed to shape regional neural timescales, whereas randomly organized networks may fail to reproduce the same multiplicity and spatial differentiation of timescales ^6,30,73^. Moreover, previous studies have linked the cortical hierarchy of INT to core–periphery organization and other macroscale anatomical gradients ^8,15^.

### Tasks modulate intrinsic neural timescales

At the same time, our results indicate that the hierarchical stability does not preclude flexible task-dependent modulation. Recent studies have reported that INT can vary systematically across behavioral states and may reflect changes in the temporal integration properties of neural circuits ^7,22^. In particular, the prolonged INT observed in some behavioral states is consistent with previous reports showing that INT increases during behavioral tasks requiring sustained attention and active engagement ^15,30,74,75^. Task-related changes have also been linked to behavioral performance: longer regional INT in the cuneus region has been associated with better mnemonic discrimination ability ^18^, suggesting that extended temporal integration may facilitate cognitive processes that require the accumulation and maintenance of information over longer timescales. Thus, flexible modulation of INT may serve as a mechanism through which the brain dynamically adapts its temporal processing to meet changing cognitive demands.

In parallel, numerous studies have demonstrated that task execution does not completely reorganize functional brain networks but instead involves constrained reconfigurations built upon the network architecture ^76–78^. Our findings further support this view. Although different tasks alter regional INT values, these changes are spatially selective rather than globally uniform. For example, social and emotional tasks tend to increase timescales in regions associated with the default mode network, whereas language and relational reasoning tasks preferentially modulate regions involved in semantic processing and cognitive control. These results suggest that task-dependent INT modulation may reflect selective adjustments of temporal integration windows within specific brain regions rather than a uniform scaling of timescales across the entire brain.

### Task-related reconfigurations of INT remain constrained by structural connectivity

Another key finding of this study is that task-related reconfigurations of INT remain constrained by structural connectivity. Previous studies have shown that the structural connectome strongly shapes the spatial organization of functional connectivity ^25,48,79^, and that network topology and hub organization play a critical role in determining large-scale brain dynamics ^80–82^. Structural constraints also apply to local dynamical properties such as intrinsic timescales. Recent resting-state literature also shows that cortical areas with greater structural strength tend to have longer timescales ^18,20,21^, and when the brain structure is altered by ischemic stroke lesions, this relationship is likely to be reshaped ^53^. Our results extend this framework to multiple tasks. Across all task conditions, the coupling strength between structural architecture and intrinsic timescales remained significantly positive, indicating that cognitive demands do not freely reconfigure neural timescales, but rather modulate temporal integration windows within an anatomically constrained dynamical space.

To further investigate the mechanisms underlying task-related changes in INT, we developed a structurally constrained multiscale model extending the influential Kinouchi-Copelli model ^37^ from a single brain region to the entire connectome. This framework is particularly well-suited for interpreting our findings within the context of brain criticality, providing a unified description of neural dynamics across scales. Theoretical studies have shown that neural systems operating near a critical point exhibit critical slowing down, characterized by slower recovery from perturbations and enhanced temporal correlations ^83^. Our simulations revealed a consistent relationship between INT and proximity to criticality across both single-region and whole-brain models. As the system approached the critical regime, INT increased, indicating long-range and long-lived correlations,. The robustness of this relationship across modeling scales suggests that the association between INT and criticality is not specific to a particular model but instead reflects a general property of neural dynamics. Moreover, recent whole-brain modeling studies have shown that large-scale brain dynamics emerge from the interplay between structural connectivity and regional heterogeneity ^28,68,84,85^, providing a natural framework for understanding how task-dependent modulations of INT arise from changes in network dynamics constrained by the underlying connectome.

Our whole-brain modeling results further suggest that the brain may not fluctuate around a single critical point but may instead operate within an extended critical regime. Theoretical studies have shown that complex networks with hierarchical and modular structure can stretch the benefits of the critical region ^56,86,87^. In such a regime, different regions of the network may simultaneously operate at different distances from criticality ^83,88–90^. Consistent with this idea, the distinct regional shifts observed across task conditions compared with the resting state may indicate regionally heterogeneous distance-to-criticality (DTC) and reveal substantial spatial heterogeneity in the direction of critical shifts. Given the spatial heterogeneity in distance to criticality (Figure 7), this finding suggests that cognitive tasks do not uniformly push the entire brain toward or away from criticality. Instead, they selectively reconfigure the dynamical states of specific brain regions or functional systems. Such nodal heterogeneity is consistent with recent perspectives from whole-brain modeling ^68,85,91,92^. Extending this conceptual framework, our findings indicate that task-related shifts in distance-to-criticality are not only spatially heterogeneous but also functionally relevant at the cognitive level.

Task-related shifts in INT at the brain region level are significantly associated with cognitive term maps derived from NeuroSynth ^43^. Regions exhibiting similar task-dependent shifts toward or away from criticality consistently correspond to specific cognitive domains, including language, emotion, social cognition, and executive control, providing a functional interpretation of the observed dynamical reconfigurations. In other words, different tasks may selectively shift functional networks closer to or farther from criticality, in line with their cognitive demands, thereby altering the temporal integration windows and coordination patterns of those networks. These findings connect structural network constraints, critical brain dynamics, and cognitive functional annotations within a unified framework. Rather than simply shifting the entire brain closer to or farther from a critical regime, task performance appears to involve selectively tuning the dynamical state of regions most relevant to the current computational demands. In this way, different brain systems may enter distinct critical operating ranges, enabling flexible configuration of neural dynamics tailored to specific cognitive domains.

### Subcortical and neuromodulatory systems in task-related intrinsic timescales

Beyond the constraints imposed by cortico-cortical structural connectivity, the spatial organization and task-dependent modulation of intrinsic neural timescales (INT) are also likely shaped by subcortical and neuromodulatory systems. In particular, the thalamus is increasingly recognized not as a passive relay, but as a central regulator of large-scale brain dynamics that influences cortical activity, interregional communication, network topology, and neuronal variability across both resting and task states ^93,94^. Through its heterogeneous and recurrent thalamocortical projections, distinct thalamic nuclei can selectively enhance task-relevant cortical interactions, sustain persistent cortical activity, and coordinate transitions between cortical representations ^95–99^. Furthermore, the complementary projection patterns of core and matrix thalamic populations align with macroscale spatiotemporal gradients of cortical organization, suggesting that thalamic architecture contributes to regional heterogeneity in cortical dynamics and intrinsic timescales ^100,101^.

The thalamus may also provide an important interface between neuromodulatory systems and cortical dynamics. Thalamic nuclei receive dense cholinergic, noradrenergic, dopaminergic and histaminergic inputs, which can alter neural excitability, gain, and the temporal properties of ongoing activity ^102–104^. More broadly, neuromodulatory regulation of neural gain has been proposed to facilitate transitions between segregated and integrated network configurations and to generate flexible, state-dependent whole-brain dynamics ^105–107^. The heterogeneous cortical distribution of neuromodulatory receptors may therefore provide a biologically plausible source of regional variation in local excitability and dynamical timescales (as seen in Figure 7). From this perspective, the relationship between structural connectivity and INT should not be viewed as a direct consequence of cortico-cortical wiring alone. Rather, it likely emerges from the interaction between structural topology, thalamocortical circuitry, and spatially heterogeneous neuromodulatory influences, which together shape the dynamical landscape underlying cognitive function.

### Limitations, and Future Directions

Despite these insights, some limitations should be acknowledged. First, INT is estimated from BOLD signal autocorrelation and may be influenced by head motion and physiological noise. Future work should incorporate more robust estimation approaches and physiological noise correction strategies ^108^. Second, our inference of criticality is largely based on modeling and indirect indicators. Future studies should combine additional criticality metrics ^83^, such as phenomenological renormalization group approaches ^88,89,109^, neuronal avalanche statistics ^110^ or spectral scaling measures ^111^, to more directly characterize the dynamical phase of brain activity across cognitive states. Third, the present study focused primarily on group-level analyses. Individual differences in structure–function coupling and critical dynamics may be substantial and could potentially relate to cognitive abilities or behavioral variability ^18,56^.

### Conclusions

In summary, the present study demonstrates that task-related changes in intrinsic neural timescales cannot be explained by structural architecture alone or by task demands in isolation. Instead, they emerge from the interaction between a largely stable cortical hierarchy of timescales and task-driven dynamical reconfigurations constrained by structural connectivity. Within this framework, distinct cognitive states can be understood as selective shifts of neural systems within a criticality landscape, with different tasks moving specific brain regions to different positions relative to the critical regime. These findings position criticality and intrinsic neural timescales as fundamental bridges connecting brain structure, neural dynamics, and cognitive function.

## Methods

All participants provided written informed consent as part of the Human Connectome Project (HCP) protocol. Data collection procedures were approved by the local Institutional Review Board. The whole-brain computational neuronal network was modeled as a random network of excitable spiking neurons. The neuronal network modeling and post-data analysis, including statistical analyses, were carried out with customized code, which is available on GitHub.

### Participants and Neuroimaging Data Acquisition

#### Participants

Data were obtained from the Human Connectome Project (HCP) S1200 public data release (March 2017). In the present study, we analyzed a subset of 339 participants who had complete resting-state fMRI and task fMRI data available for analysis ^112^. Following quality control and the exclusion of participants with missing practice-related data, 326 participants were retained for the practice-related analyses. The HCP dataset includes seven task paradigms that probe different cognitive domains: working memory, social cognition, motor function, language processing, gambling-based decision-making, emotional processing, and relational reasoning. Only participants with usable data across these modalities were included in the final sample. No statistical methods were used to predetermine sample sizes, but our sample sizes are similar to those reported in previous publications using the full HCP dataset.

#### Functional MRI Acquisition

Functional MRI data were acquired using a 3 Tesla Siemens Connectome Skyra scanner with a 32-channel head coil. In addition to resting-state scans, the HCP protocol includes multiple task-based fMRI paradigms designed to assess different cognitive functions. For resting-state fMRI, participants were scanned for approximately 15 minutes while maintaining visual fixation on a projected crosshair presented on a dark background with their eyes open. Task-based fMRI data included seven paradigms: working memory, social cognition, motor, language, gambling, emotion, and relational processing. Detailed descriptions of task paradigms and acquisition parameters are provided in the official HCP documentation (http://www.humanconnectome.org).

#### The HCP repository of seven tasks

The HCP task repository comprises seven tasks: working memory, motor, gambling, language, social cognition, emotion, and relational processing. Detailed descriptions of these paradigms are provided in the Table 1. These tasks were developed to sample a wide range of cognitive processes and associated neural systems. Specifically, they target several major functional domains, including sensory–motor processing (visual, motion, somatosensory, and motor functions), cognitive control and working memory, decision-making processes, category-specific representations, language comprehension, relational reasoning, social cognition, and emotional processing.

All participants in the HCP dataset completed the full set of tasks across two scanning sessions. The first session included the working memory, gambling, and motor tasks, whereas the second session comprised the language, social cognition, relational processing, and emotion tasks. In addition to the main cohort, a subset of 45 participants underwent a retest session, allowing assessment of test–retest reliability for the task paradigms.

#### Diffusion MRI Acquisition

Diffusion MRI data used for structural connectivity analysis were obtained from the HCP database. Diffusion-weighted images were acquired using a multi-shell diffusion scheme with three diffusion weightings (b = 1000, 2000, and 3000 s/mm²). The standard acquisition protocol consisted of six runs totaling approximately 59 minutes. Further details regarding the diffusion imaging protocol are available in the HCP documentation.

### MRI Data Preprocessing and Brain Parcellation

#### Brain parcellation

To define cortical regions for analysis, we primarily used the Mindboggle-modified Desikan–Killiany atlas, which partitions the cortex into 62 regions (31 per hemisphere) ^113,114^. This parcellation provides a coarse but widely used representation of cortical organization and has been extensively applied in studies of large-scale brain dynamics. All analyses reported in the main text were performed using this DK62 cortical parcellation. To evaluate the robustness of our findings across spatial scales, we additionally repeated the analyses using the Glasser atlas ^42^, which divides the cortex into 360 regions (180 per hemisphere, namely Glasser360). Results obtained with this higher-resolution parcellation are reported in the Supplementary Information. Both parcellation schemes were defined in the HCP CIFTI grayordinates standard space, enabling direct extraction of regional signals from the minimally preprocessed functional data.

#### fMRI preprocessing

Resting-state and task fMRI data were obtained from the Human Connectome Project minimal preprocessing pipeline, which integrates tools from FSL, FreeSurfer, and Connectome Workbench ^115,116^. The preprocessing procedures include correction for spatial distortions, head motion, and gradient-related artifacts, followed by intensity normalization and bias-field correction. Functional images were aligned to the individual structural scans and transformed to MNI space ^117^.

To further reduce non-neuronal noise, structured artifacts were identified using independent component analysis and removed with the ICA-FIX denoising procedure ^118,119^. Motion-related regressors were also included to minimize residual head-motion effects. The preprocessed data were provided in CIFTI grayordinates format, which combines cortical surface and subcortical volumetric representations. Regional BOLD timeseries were then extracted based on the selected parcellation. For each cortical parcel, the mean signal across all grayordinates within the parcel was computed ^120^. The resulting timeseries were temporally filtered using a second-order Butterworth band-pass filter (0.008–0.08 Hz) to retain the low-frequency fluctuations associated with large-scale functional dynamics ^121^.

#### Diffusion MRI preprocessing and connectome

The provided participants have high-quality whole-brain diffusion imaging with a total acquisition time of approximately 59 minutes. Diffusion data were processed using pipelines implemented in the Lead-DBS framework ^122^, following previously established procedures ^123–129^. Fiber orientation distributions were reconstructed using generalized q-sampling imaging as implemented in DSI Studio. Whole-brain tractography was then performed to reconstruct white-matter pathways. For each participant, a large set of streamlines was generated within the white-matter mask and subsequently transformed to Montreal Neurological Institute (MNI) standard space. These reconstructed fiber pathways were then used to estimate the structural connectome. The resulting tractography-based connectivity matrices were mapped to the cortical parcellation introduced above to construct the structural connectivity (SC) matrices (denoted as **SC**), and they were used to guide the construction of the whole-brain neuronal network model (see section "Whole-brain Neuronal Network Modeling"). The SC strength was defined as the weighted degree of the structural connectome, where 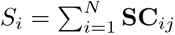, *N* corresponds to the number of regions defined by the parcellation. Following previous work ^130^, connection strengths were normalized by the mean value across nodes, i.e.,*S_i_* = *S_i_/ < S_j_ >*.

### Neuronal network modeling

#### Single regional neuronal network

Each functional brain region was modeled as a local neuronal network composed of excitable units Figure 1**F**. Specifically, within each region, we constructed an Erdő s–Rényi undirected random graph consisting of *n* = 10,000 neurons with an average degree of *K* = 10. In this local network, nodes represented individual excitable neurons, and edges represented local neuronal interactions. The Erdő s–Rényi topology was used to capture sparse and random local connectivity while preserving a controlled average degree across regions.

#### Whole-brain neuronal network

We modeled the brain as a whole-brain network of interconnected functional regions. At the macroscale, each node represented a functional brain region defined by the cortical parcellation scheme Figure 1**G**. Each brain region was further modeled at the microscale as a large Erdös–Rényi undirected random graph representing the local neuronal network within that region. Specifically, each functional region *i* was represented by a local neuronal network, consisting of *n* = 10, 000 excitable neurons organized as an Erdös–Rényi random graph with average degree *K* = 9.65. Thus, each region was modeled as a large-scale local network capturing the intrinsic neuronal interactions within that region.

At the macroscale, interregional coupling was constrained by the empirical structural connectome derived from diffusion MRI tractography. Specifically, structural connectivity between regions was quantified by the number of streamlines reconstructed between each pair of regions, yielding a structural connectivity matrix [*SC_ij_*], where *S_ij_* denotes the structural connection strength between regions *j* and *j* (Figure 1**G**).

To determine the number of long-range connections between regions, the structural connectivity matrix was normalized as

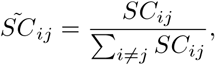

such that 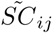 represents the relative proportion of interregional connectivity between regions *i* and *j*. The average interregional degree between regions *i* and *j* was then defined as

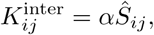

where *α* is a scaling factor. The scaling factor *α* was chosen such that the maximally connected region had a total degree close to *K*_total_ = 10. Given a fixed intraregional degree of *K*_intra_, the target maximum interregional degree was 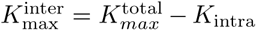. Therefore, *α* was defined as

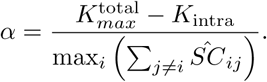

Given that each region contained *n* neurons, the number of interregional neuron-to-neuron connections between regions *i* and *j* was defined as

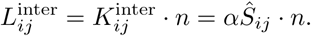

For each pair of connected regions, the 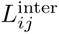 long-range links were implemented by randomly selecting neurons from regions *i* and *j* and establishing neuron-to-neuron connections between them. This procedure generated a heterogeneous whole-brain network in which the total number of connections associated with region *i* was given by

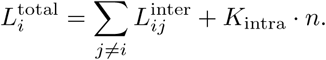

Thus, the large-scale organization of the whole-brain model was constrained by the empirical structural connectome, while each region retained its own local random network architecture.

#### Neuronal Dynamics

For a given functional region *i*, the neuron population consisted of *n* coupled excitable neurons. Neuronal interactions within and between regions were described by the network topology defined above. The network degree of region i is set by 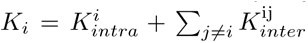 and average network degree is given by 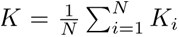. The Kinouchi & Copelli model ^37^ was then used to simulate the dynamics. Each neuron was modeled as a cyclic cellular automaton evolving in discrete time over three states: susceptible (*x* = 0), active (*x* = 1), and refractory (*x >* 1). Neuronal states were updated synchronously at each discrete time step of duration *δ_t_* = 1 ms. When a neuron was in the susceptible state at time *t*, it could be excited in one of two ways: (i) by an active neighboring neuron with transmission probability *λ* = 0.1, or (ii) by an external stimulus modeled as a Poisson process with rate *r* = 10*^−^*^5^. The corresponding activation probability of the external input was *h* = 1 *−* exp(*−r*). Thus, the joint excitation probability for a susceptible neuron was given by *q* = 1 *−* (1 *− h*)(1 *− λ*). An active neuron at time *t* will go to the refractory state at time *t* + 1 and remain there for 8 ms. After that, the neuron will enter a susceptible state. See the flowchart in Figure 1 for a schematic illustration of the neuronal dynamics.

#### Branching Ratio and Distance to Criticality

The density of active neurons, *F* 0*_i_*(*t*), was used to characterize the activity of the whole-brain neuronal network as well as the neuronal network in region *i* and to compute its intrinsic neural timescale.

Given that the total average degree of region *i* was 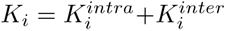. The branching ratio of region *i* was defined as 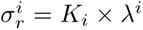, which represents the average number of activations generated by one active neuron at the next time step ^37^. Critical branching occurs when *σ_i_*= 1. For *σ_i_ <* 1, activity decays over time, indicating a subcritical regime. For *σ_i_ >* 1, activity grows over time, indicating a supercritical regime ^131^. Thus, the distance to criticality for region *i* in the subcritical regime was quantified as 1 *− σ_i_*.

At the whole-brain level, the global average branching ratio across all regions can be written as:

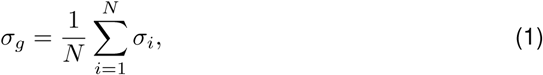

where *N* is the number of functional regions. Correspondingly, the mean distance to criticality of the whole-brain network is 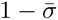.

### Autocorrelation Function and Intrinsic Neural Timescales

With the neural activity of brain functional networks and neuronal networks, intrinsic neural timescales were computed based on the autocorrelation function (ACF) of their timeseries, where

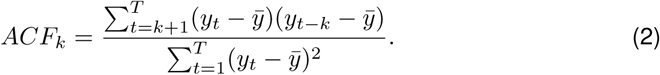

*y* denotes the timeseries of neural activity, *y̅* is the mean value across time points, *t* is the time step. The intrinsic timescales were then computed as the area under the curve (AUC) of the initial ACF curve until it reaches a zero value:

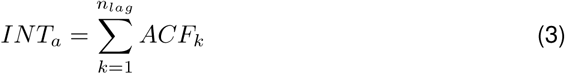

With BOLD timeseries and the modeling dynamics, intrinsic timescales for functional and modeled whole-brain neuronal networks were generated. The intrinsic timescales for specific large-scale functional networks were averaged to yield the network-level intrinsic timescale. The intrinsic timescales of large-scale functional networks were obtained to investigate the hierarchy of INT across the functional network.

### Whole-brain INT and Neurosynth Associations

To characterize task-related changes in intrinsic neural timescales (INT), we first computed a whole-brain INT map for each fMRI condition. For each task condition *c*, the task-related INT difference map was defined as

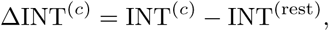

where INT^(*c*)^ and INT^(rest)^ denote the regional INT maps for the task and restingstate conditions, respectively. These task–rest difference maps were used to quantify the spatial pattern of task-related INT reconfiguration across the brain.

To investigate the cognitive relevance of these spatial patterns, we used Neurosynth ^43^, an automated meta-analytic platform that provides large-scale association maps for cognitive terms. Association maps for 89 cognitive terms were obtained and parcellated using the same atlas as in the present study. For each task condition, we correlated the corresponding ΔINT^(*c*)^ map with each parcellated Neurosynth association map. The absolute correlation coefficient was used to quantify the strength of association between task-related INT reconfiguration and each cognitive term.

To account for spatial autocorrelation in brain maps, we performed Spin Test analyses ^132^ using the BrainSMASH Python library ^133^. This procedure was applied to all correlation analyses involving cortical surface data. Spatially constrained null maps were generated while preserving the spatial autocorrelation structure of the empirical maps. We used 10,000 permutations to generate null distributions of correlation values, and empirical correlations were compared against these null distributions to obtain p-values corrected for spatial autocorrelation. False discovery rate (FDR) correction was further applied across all comparisons.

### SC-INT coupling

To examine whether anatomical connectivity constrains intrinsic neural timescales (INT), we quantified the coupling between regional structural connectivity strength and regional INT values across rest and task states. For each cortical region, nodal SC strength was calculated as the sum of its weighted connections to all other regions:

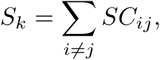

where *SC_ij_*denotes the structural connectivity weight between regions *i* and *j*.

For each participant and each condition, SC-INT coupling was defined as the spatial correlation between regional SC nodal strength and regional INT values:

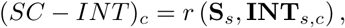

where **S***_s_* is the vector of regional SC strength for participant *s*, and **INT***_s,c_* is the vector of regional INT values for participant *s* under condition *c*. This analysis was performed separately for rest and each task condition. The resulting subject-level SC–INT coupling values were used to assess whether structure–timescale coupling varied across cognitive states.

In addition to subject-level coupling, we examined group-level node-wise SC–INT relationships. For each condition, regional SC strength and INT values were averaged across participants, and Pearson spatial correlations were computed across cortical regions. These node-wise spatial correlations were used to characterize the extent to which regions with stronger structural connectivity exhibited longer intrinsic neural timescales at the group level. The SC–INT coupling analysis was repeated using both the DK62 and Glasser360 parcellations to evaluate robustness across spatial resolutions.

#### Similarity and distance

To quantify task-related modulation of intrinsic neural timescales, we computed regional INT difference maps for each task condition relative to rest. For each task *i* and cortical region *k*, the INT difference was defined as:

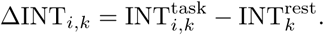

Positive values indicate regions with longer INT during task relative to rest, whereas negative values indicate regions with shorter INT during task relative to rest.

To assess whether different tasks induced similar or distinct spatial patterns of INT modulation, each task-specific INT difference map was vectorized across cortical regions and compared pairwise across task conditions. We first quantified spatial similarity using Pearson correlation:

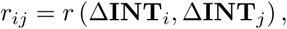

where Δ**INT***_i_* and Δ**INT***_j_* denote the vectorized INT difference maps for tasks *i* and *j*. Spatial dissimilarity was then quantified using correlation distance:

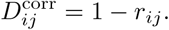

Smaller values indicate more similar task-induced INT modulation patterns, whereas larger values indicate more distinct spatial configurations.

#### Statistics and Reproducibility

Statistical analyses were performed using standard parametric and non-parametric tests where appropriate. Group-level differences in INT across rest and task conditions were assessed using repeated-measures analysis of variance (rmANOVA). When the sphericity assumption was violated, the Greenhouse-Geisser correction was applied to adjust the degrees of freedom. For significant rmANOVA results, *post hoc* pairwise comparisons between rest and each task condition were performed using paired-samples *t*-tests. Multiple comparisons were controlled using the Benjamini–Hochberg false discovery rate procedure, with statistical significance defined as *p_F DR_ <* 0.05. This correction was applied to post hoc task comparisons, node-wise spatial correlation analyses, and cognitive decoding results where appropriate.

To assess the preservation of temporal hierarchy across cognitive states, regional INT values were averaged within large-scale functional networks. Networks were ranked according to their mean INT values under each condition, and the similarity between task-specific and resting state rank orders was quantified using Spearman rank correlation. Correlations between SC strength and INT values were assessed using Pearson correlation for spatial node-wise analyses and subject-level SC–INT coupling analyses.

For computational modeling analyses, simulations were repeated across 50 independent trials for each value of *σ*. Optimal *σ* values were estimated separately for each trial, and the mean and variability across trials were reported. Model reproducibility was further evaluated by comparing condition-level empirical BOLD-derived INT values with model-derived INT values at the optimal *σ* using Spearman correlation across rest and task conditions.

All statistical analyses, SC-INT coupling analyses, similarity, spin permutation tests, cognitive decoding analyses, and computational neuronal network simulations were performed using customized code. The code is made available online, as detailed in the “Code availability” section.

## Data Availability

All data used in this study are publicly available, including functional MRI (fMRI), diffusion MRI, brain atlases, modeling dynamics, and Neurosynth-derived functional association maps. The fMRI and dMRI datasets were obtained from publicly accessible neuroimaging repositories. The two brain atlases used for cortical parcellation and model construction are publicly available from their original sources. The Neurosynth-derived maps are available through the Neurosynth platform. The modeling dynamics generated and analyzed in this study are publicly available upon publication. All necessary data are available in the GitHub repository associated with this study: https://github.com/kaichaowuu/ Structure-Constrained_Intrinsic_Timescales_for_Rest_and_Across_Tasks.

## Code Availability

The code for neuronal networks modeling, post-processing analyses including the intrinsic timescale estimation and statistical analyses, as well as the result visualization, are all available here GitHub.

## Acknowledgements

This work was supported by the Australian Research Council, Future Fellowship (FT200100942), the Juan De La Cierva (JDC2024-055992-I) and Ramón y Cajal Fellowship (RYC2022-035106-I) from FSE/Agencia Estatal de Investigación (AEI), and funded by the Spanish Ministerio de Ciencia, Innovación y Universidades (MICI-U/AEI/10.13039/501100011033) through the María de Maeztu project (CEX2021-001164-M). The funder played no role in study design, data collection, analysis and interpretation of data, or the writing of this manuscript.

## Author Contributions

KC.W: Conceptualization, Data curation, Software, Investigation, Formal analysis, Validation, Visualization, Writing – original draft, Writing – review & editing. RR.P: Writing – review & editing. R.H: Writing – review & editing. C.M: Writing – review & editing. P.M: Writing – review & editing. LL.G: Conceptualization, Investigation, Methodology, Supervision, Funding acquisition, Writing – original draft, Writing – review & editing.

## Supplementary information

The supplemental materials and data are available in the attached Supplementary Materials.

