## supplementray for "Structure-Constrained Intrinsic Timescales Across Tasks"

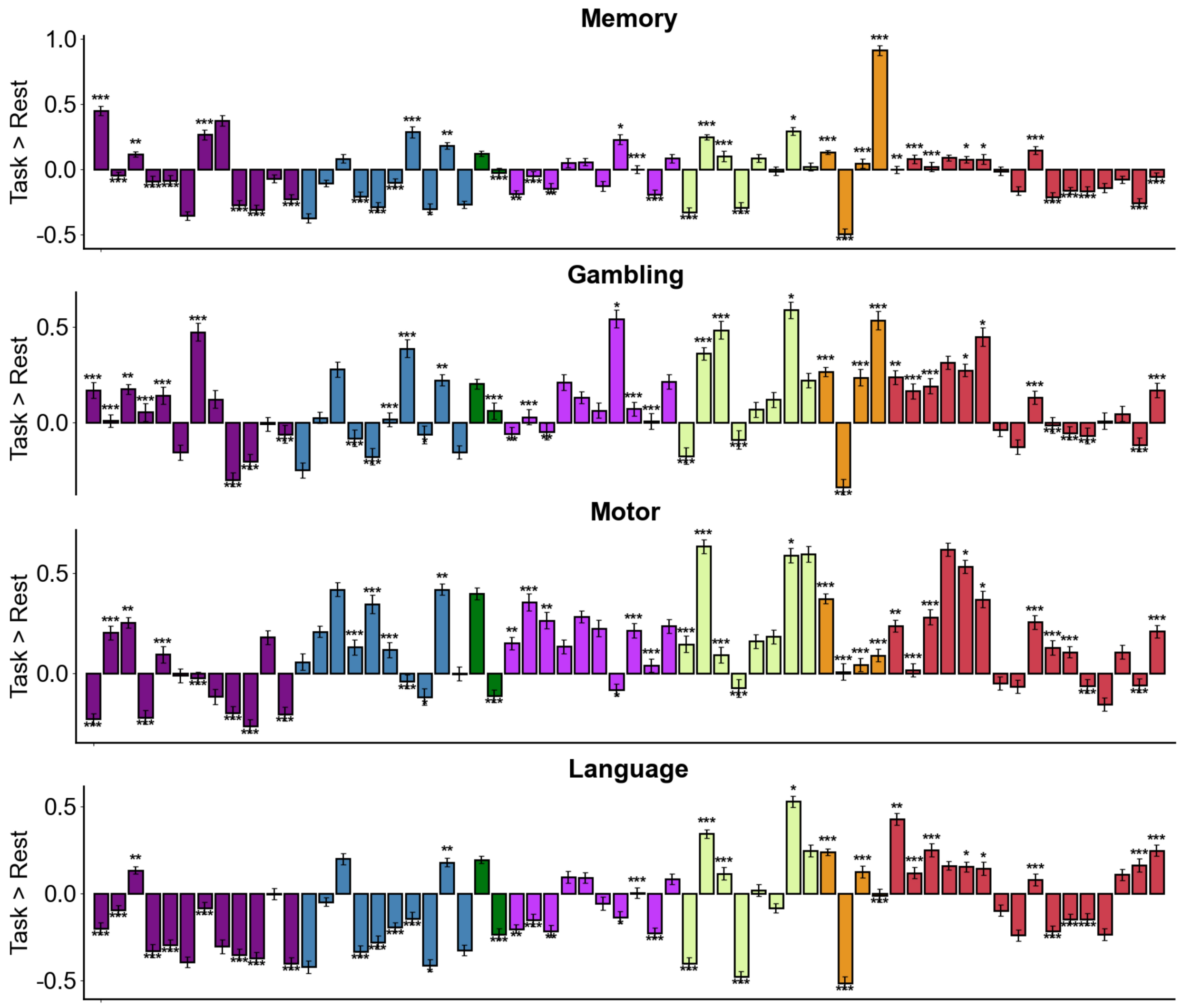

Memory
Gambling
Motor
Language
Task > Rest
1.0
0.5
0.0
-0.5

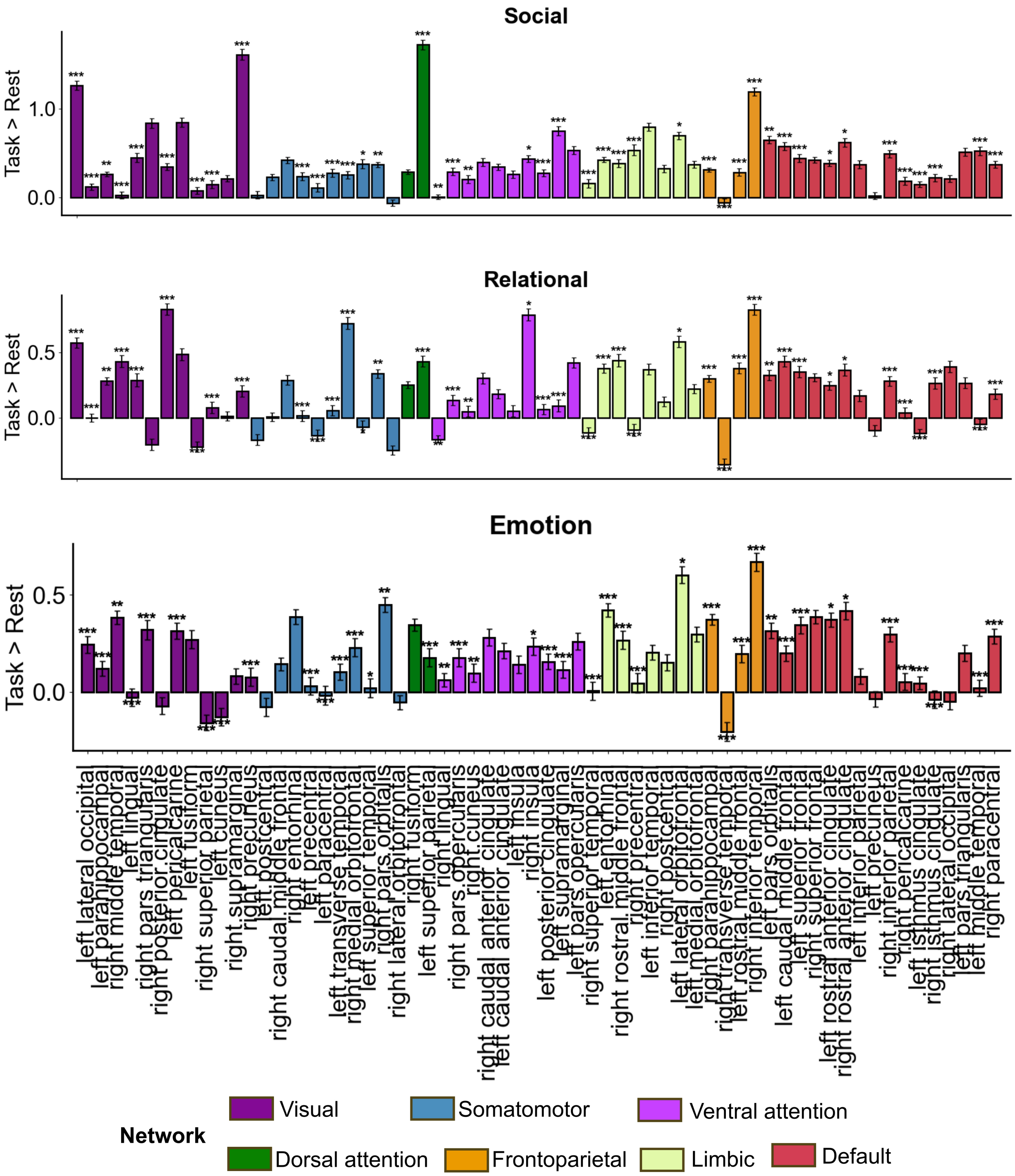


**Supplementary Figure 1 – Regional task-induced changes in intrinsic neural timescales relative to rest.** Bar plots show the regional INT differences between each task condition and the resting state for the Memory, Gambling, Motor, Language, Social, Relational, and Emotion tasks. The y-axis indicates the direction and magnitude of task-related INT modulation, with positive values indicating longer INT during task relative to rest and negative values indicating shorter INT during task relative to rest. Each bar represents one cortical region, and colors denote seven large-scale functional networks. Error bars indicate variability across participants. Asterisks indicate regions showing significant task–rest differences after multiple-comparison correction. Overall, task-induced INT changes were spatially heterogeneous, with both increases and decreases observed across cortical regions and task domains.

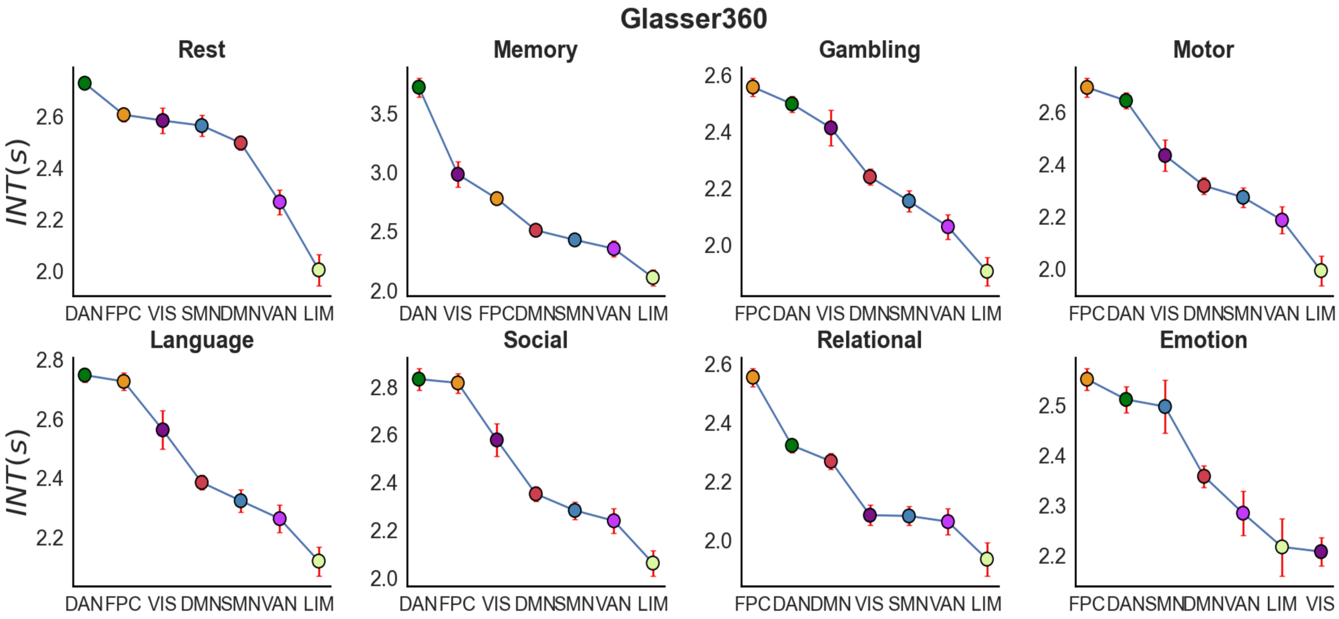


**Supplementary Figure 2 – Stable hierarchy of INTs is also supported by the Glasser360 parcellation.** Across tasks, the correlation coefficients between the hierarchy rank of task and rest ranged from $rho = -0.32$ to $rho = 0.11$ with all comparisons being non-significant ($p_{FDR} > 0.33$), demonstrating that the brain temporal hierarchy was largely preserved across tasks. Functional network abbreviations are as follows: dorsal attention (DAN), frontoparietal control (FPC), somatomotor (SMN), visual (VIS), default (DMN), ventral attention (VAN), and limbic (LIM).

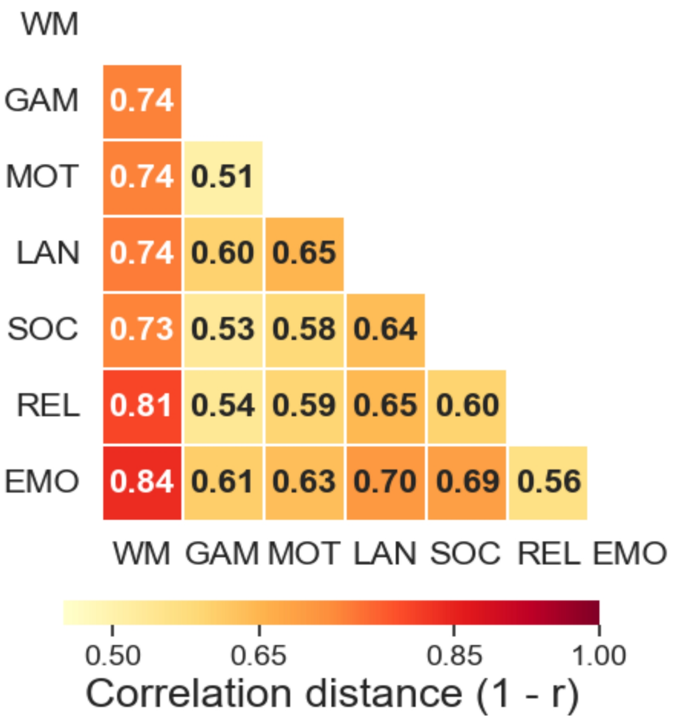


**Supplementary Figure 3 – Pairwise spatial distance between task-specific INT difference maps for the Glasser360 parcellation.** Each INT difference map was computed as task minus rest and vectorized across cortical regions. Working Memory (WM), Gambling (GAM), Motor (MOT), Language (LAN), Social (SOC), Relational (REL), and Emotion (EMO). The distance between two tasks was defined as correlation distance, $1-r$, where $r$ denotes the Pearson spatial correlation between their INT difference maps. Smaller values indicate more similar spatial patterns of task-induced INT modulation, whereas larger values indicate more distinct task-related INT changes. The smallest distance was observed between Gambling and Motor ($D = 0.51$), whereas the largest distance was observed between Working Memory and Emotion ($D = 0.84$). The color bar indicates correlation distance $(1-r)$, with lighter colors representing smaller distances and more similar INT difference maps, and darker red colors representing larger distances and more distinct INT difference maps.

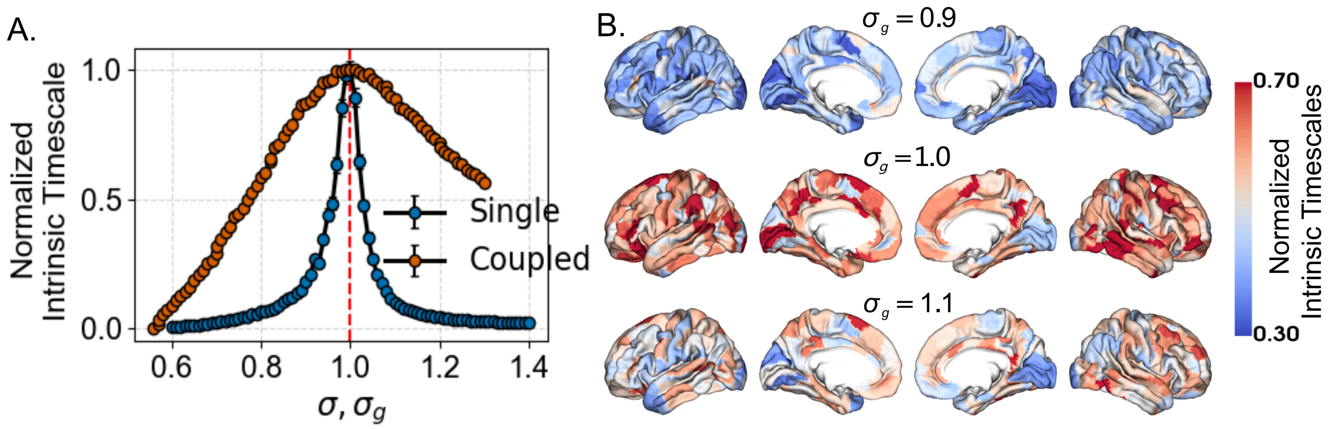


**Supplementary Figure 4 – Modeling reveals intrinsic timescales and their relationship with structural connectivity across dynamical regimes for the Glasser360 parcellation.** Intrinsic timescales as a function of the control parameter $\sigma$ for single region (blue) and $\sigma_g$ for whole-brain coupled (red) networks. In the isolated single region, intrinsic timescales exhibit a sharp peak at $\sigma \approx 1$. The coupled network also peaks at $\sigma_g \approx 1$, whereas it shows a broader peak, indicating an extended near-critical regime of elevated timescales. **(B)** Spatial maps of intrinsic timescales show regime-specific heterogeneity across regions. The color bar indicates the normalized intrinsic timescale.
